# BSCAMPP: Batch Scaled Phylogenetic Placement on Large Trees

**DOI:** 10.1101/2022.10.26.513936

**Authors:** Eleanor Wedell, Chengze Shen, Tandy Warnow

## Abstract

Phylogenetic placement, the problem of placing sequences into phylogenetic trees, has been limited either by the number of sequences placed in a single run or by the size of the placement tree. The most accurate scalable phylogenetic placement method with respect to the number of query sequences placed, EPA-ng, has a runtime that scales sublinearly to the number of query sequences. However, larger phylogenetic trees cause an increase in EPA-ng’s memory usage, limiting the method to placement trees of up to 10,000 sequences. Our recently designed SCAMPP framework has been shown to scale EPA-ng to larger placement trees of up to 200,000 sequences by building a subtree for the placement of each query sequence. The approach of SCAMPP does not take advantage of EPA-ng’s parallel efficiency since it only places a single query for each run of EPA-ng. Here we present BATCH-SCAMPP, a new technique that overcomes this barrier and enables EPA-ng and other phylogenetic placement methods to scale to ultra-large backbone trees and many query sequences. BATCH-SCAMPP is freely available at https://github.com/ewedell/BSCAMPP_code.

## 1 Introduction

Phylogenetic placement is the problem of placing one or more query sequences into a phylogenetic “backbone” tree, which may be a maximum likelihood tree on a multiple sequence alignment for a single gene, a taxonomy with leaves labeled by sequences for a single gene [1], or a species tree [2]. When the backbone tree is a tree estimated on a single gene, the most accurate techniques for phylogenetic placement are likelihood-based, and can be computationally intensive when the backbone trees are large [3]. Phylogenetic placement into gene trees occurs when updating existing gene trees with newly observed sequences, but can also be applied in the “bulk” context, where many sequences are placed at the same time into the backbone tree. For example, it can be applied when seeking to taxonomically characterize shotgun sequencing reads generated for an environmental sample in metagenomic analysis [1, 4, 5].

Thus, phylogenetic placement is a general tool with applications in both incremental tree building and taxon identification and abundance profiling in microbiome analysis [1, 4, 6–9]. In these analyses, however, the number of sequences placed into the backbone tree can be very large, for example, when attempting to place all the shotgun sequencing reads in the sample into a taxonomic tree. Even approaches that only place a subset of the reads that map to marker genes (i.e., genes that are considered single-copy and universal) can require placement of thousands to tens of thousands of reads into the taxonomic tree for that marker gene [4]. Moreover, such approaches can have additional computational challenges as the taxonomic tree they are placing into may be very large, with perhaps many thousands or hundreds of thousands of leaves.

Thus, two major aspects impact the runtime (and potentially accuracy) of phylogenetic placement methods: the size of the backbone tree and the number of sequences being placed into the tree. Since many studies have shown the improvement in accuracy for abundance profiling, phylogenetic tree estimation, etc., when placing into large backbone trees (e.g., [1]), the importance of using phylogenetic placement methods that can run on large datasets is well established.

However, not all phylogenetic placement methods can scale to large backbone trees (and many methods are computationally intensive on large trees) [10, 11], which has led to the development of methods, such as APPLES-2 [12], which use distances to place into large trees. There are also methods for phylogenetic placement that are alignment-free, such as RAPPAS [13] and App-SpaM [14]. These methods are potentially faster and more scalable than likelihood-based analyses, such as pplacer [15] and EPA-ng [5]. In general, both the number of query sequences (i.e., sequences that need to be placed into trees) and the size of the backbone tree represent challenges for phylogenetic placement methods, especially when seeking to use likelihood-based placement methods.

Previously we introduced the SCAMPP framework [10] to enable both pplacer and EPA-ng to perform phylogenetic placement into ultra-large backbone trees, and we demonstrated its utility for placing into backbone trees with up to 200,000 sequences. By using maximum likelihood methods pplacer or EPA-ng within the SCAMPP framework, the resulting placements are more accurate than with APPLES-2, with the most notable accuracy improvement for fragmentary sequences, and are computationally similar for single query sequence placement [10]. However, SCAMPP was designed to incrementally update a large tree, one query sequence at a time, and was not optimized for the other uses of phylogenetic placement, where batch placement of many sequencing reads is required.

EPA-ng is optimized for scaling the number of query sequences and is capable of placing millions of sequences into phylogenetic trees of up to a few thousand sequences [5]. EPA-ng performs this batch placement very efficiently, achieving sublinear runtime in the number of query sequences (see Figure 2 from [12]). However, other studies have shown that EPA-ng has a large memory requirement that depends on the size of the backbone tree; hence, the backbone trees used with EPA-ng are typically limited to a few thousand sequences.

Here we introduce BATCH-SCAMPP, a technique that improves scalability in both dimensions: the number of query sequences being placed into the backbone tree and the size of the backbone tree. Furthermore, BATCH-SCAMPP is specifically designed to improve EPA-ng’s scalability to large backbone trees.

BATCH-SCAMPP is based on SCAMPP, our prior method for scaling phylogenetic placement methods to large backbone trees. However, by its design, SCAMPP is not suitable for batch placement. Specifically, SCAMPP operates one query sequence at a time. Given the query sequence, SCAMPP finds and extracts a single (small) placement subtree out of the large backbone tree and then uses the selected phylogenetic placement method to place the query sequence into that placement subtree. Then, by superimposing the placement tree into the backbone tree and considering branch lengths, SCAMPP can identify the correct location within the original backbone tree to add the query sequence. While this approach enables phylogenetic placement methods to scale to ultra-large backbone trees, the fact that a different subtree is selected for each query sequence makes this approach ineffective for batch placement. Hence,

BATCH-SCAMPP uses a substantially modified design in order to be able to take advantage of EPA-ng’s ability to place many query sequences efficiently. The BATCH-SCAMPP method operates by allowing the input set of query sequences to suggest and then vote on placement subtrees, thus enabling many query sequences to select the same placement subtree. We pair BATCH-SCAMPP with EPA-ng to explore the capability of this approach for scaling to many query sequences. We show that this combination of techniques (which we call BSCAMPP+EPA-ng, or BSCAMPP(e)) not only provides high accuracy and scalability to large backbone trees, matching that of SCAMPP used with EPA-ng (i.e., SCAMPP(e)), but also achieves the goal of scaling sublinearly in the number of query sequences. Moreover, it is much more scalable than EPA-ng and faster than SCAMPP+EPA-ng: when placing 10,000 sequences into a backbone tree of 50,000 leaves, EPA-ng could not to run due to memory issues, SCAMPP+EPA-ng required 1421 minutes, and BSCAMPP(e) placed all sequences in 7 minutes (all given the same computational resources, see Table 2).

The rest of the paper is organized as follows. We begin in Section 2 with preliminary experiments evaluating prior phylogenetic placement methods, motivating the need for a novel divide-and-conquer strategy to improve the scalability of EPA-ng. We then present our design of BSCAMPP(e) in Section 3. The experimental study where we design and evaluate BSCAMPP is described in Section 4, and the results are presented in Section 5. We provide a discussion of results in Section 6, and we finish with conclusions in Section 7. Due to space constraints, some of the results are provided in the Supplementary Materials.

## 2 Preliminary Experiment

We performed a preliminary experiment that studied EPA-ng, both in terms of its accuracy and its scalability (with the number of query sequences or the size of the backbone tree). The results of this experiment motivate the goals in developing a technique to enable EPA-ng to scale to large backbone trees, and also provide some insight into how to design such a strategy. For this experiment, we used two simulated datasets from prior studies evaluating phylogenetic placement methods: two subsets of RNASim [16] with up to 180,000 sequences in the backbone tree (placing at most 20,000 query sequences) and nt78 [17] with 68,132 sequences in the backbone tree (placing 10,000 query sequences). We explore phylogenetic placement error using the “delta error” (see Section 4.7) and runtime.

### 2.1 Experiment 0: Understanding EPA-ng

Here we focus on EPA-ng for placing short sequences (the metagenomics context, see Supplementary Materials for results when placing also full-length sequences). We extracted a subset of the RNASim training dataset (see Section 4.4) for this experiment. The subset is a clade with 10,200 leaves of the training RNASim dataset, from which we randomly selected 200 leaves as queries. For the remaining 10,000 leaves, we randomly selected from 1000 to 10,000 leaves to form the backbone tree. For the queries, we made them into fragments ranging from 10%-length and 50%-length. All queries were placed into the backbone trees using EPA-ng in one of two modes: “batch”, where we place all 200 queries all at once (a feature allowed by EPA-ng), and “one-by-one”, where we place each query independently by running EPA-ng on each query. We show placement error (using delta error, see Section 4.7 for definition) on various backbone tree sizes and placement strategies (batch vs. one-by-one). Results for 10%-through 50%-length are shown in Figure S2 in the Supplement.

For both query sequence lengths, accuracy improves for one-by-one placement as the backbone size increases, indicating the beneficial impact of taxon sampling. Interestingly, this is not true for batch placement. For batch placement, phylogenetic placement accuracy is very much impacted by query length, but very fragmentary sequences are only placed well on small backbone trees (up to 2,000 leaves).

Since our interest is in placing short sequences (corresponding to read placement), we focus now on the results shown for queries of 10%-length. For these very short query sequences, we see a jump in delta error for “batch” from backbone tree size of 2000 to 5000 and up. Thus, high accuracy is only obtained when placing into smaller trees. Thus, EPA-ng performs very differently between the batch mode (the case where it is able to obtain a runtime advantage, according to prior studies) and the one-by-one mode. When the backbone tree is small enough then batch mode can be very accurate, but on larger backbone trees the one-by-one mode is more accurate for placing fragmentary sequences. The key lesson is that if we wish to obtain good accuracy in placing fragmentary sequences, we need to either use EPA-ng in one-by-one mode or keep the backbone tree to at most 2000 sequences.

We next performed an experiment to understand the impact of varying the number of query sequences placed with EPA-ng and pplacer. Query sequences are all made fragmentary to 10% of the full sequence length. We show results on a 1000-leaf backbone tree for various numbers of query sequences (Figure S1 in the Supplement). Note that EPA-ng takes less time to place 100 and 1000 sequences than it does to place 10 sequences, and more generally that its runtime grows sublinearly in the number of query sequences. Furthermore, neither its peak memory usage nor the delta error notably increases when placing more queries. In contrast, pplacer’s runtime grows much more quickly than EPA-ng’s.

These trends show that EPA-ng is fast when run in batch mode (instead of one-by-one), but only when placing into relatively small trees. Moreover, when placing into relatively small trees (i.e., less than 5000 leaves) using batch mode, its runtime is sublinear in the number of query sequences. These two observations motivate the development of BATCH-SCAMPP, the subject of the next section.

## 3 BATCH-SCAMPP

BATCH-SCAMPP (which we will call BSCAMPP for short) was designed with the goal of developing a divide-and-conquer strategy that allowed EPA-ng to run on datasets with large backbone trees, however, this approach could be used with any phylogenetic placement method. We assume that we have an input backbone tree *T* and its associated multiple sequence alignment of the sequences at the leaves of the tree, and a set *Q* of query sequences that we will place into the tree *T*.

At a high level, our approach has four stages: Stage 1: the sequences in *Q* vote for their top *v* placement subtrees, each of bounded size *B* (both parameters specified by the user); Stage 2: We construct a set 𝒯 of placement subtrees using the votes and we assign each query in *Q* to one of the placement subtrees; Stage 3: We allow reassignments of query sequences to subtrees; and Stage 4: we run EPA-ng on the placement subtrees to add the assigned query sequences into the subtrees, and then use branch lengths to find appropriate positions in the backbone tree *T*.

This four-stage approach is an elaboration on the SCAMPP technique, except that in SCAMPP, each query sequence picks a single placement subtree, and it is completely feasible that there will be as many placement trees as there are query sequences.

In a collection of initial experiments (all on the same training data), we explored four different variants of this four-stage approach (described in S2.1 in the Supplementary Materials). These variants differ mainly in Stage 2, which uses the votes to determine the subtree to assign each query sequence. In comparing the different strategies, we optimized both accuracy and computational effort, and selected variant 4, which had the best combined accuracy and speed. This variant requires that all queries can be placed into a subtree containing its most similar sequence (by Hamming distance) and that once our greedy subtree selection has finished queries can be reassigned to any of the selected subtrees which minimize the queries total distance to other all leaves in the tree. We refer to the selected variant as BATCH-SCAMPP or BSCAMPP, noting that it has two algorithmic parameters (subtree size *B* and the number *v* of votes).

**Algorithm BSCAMPP(e)**

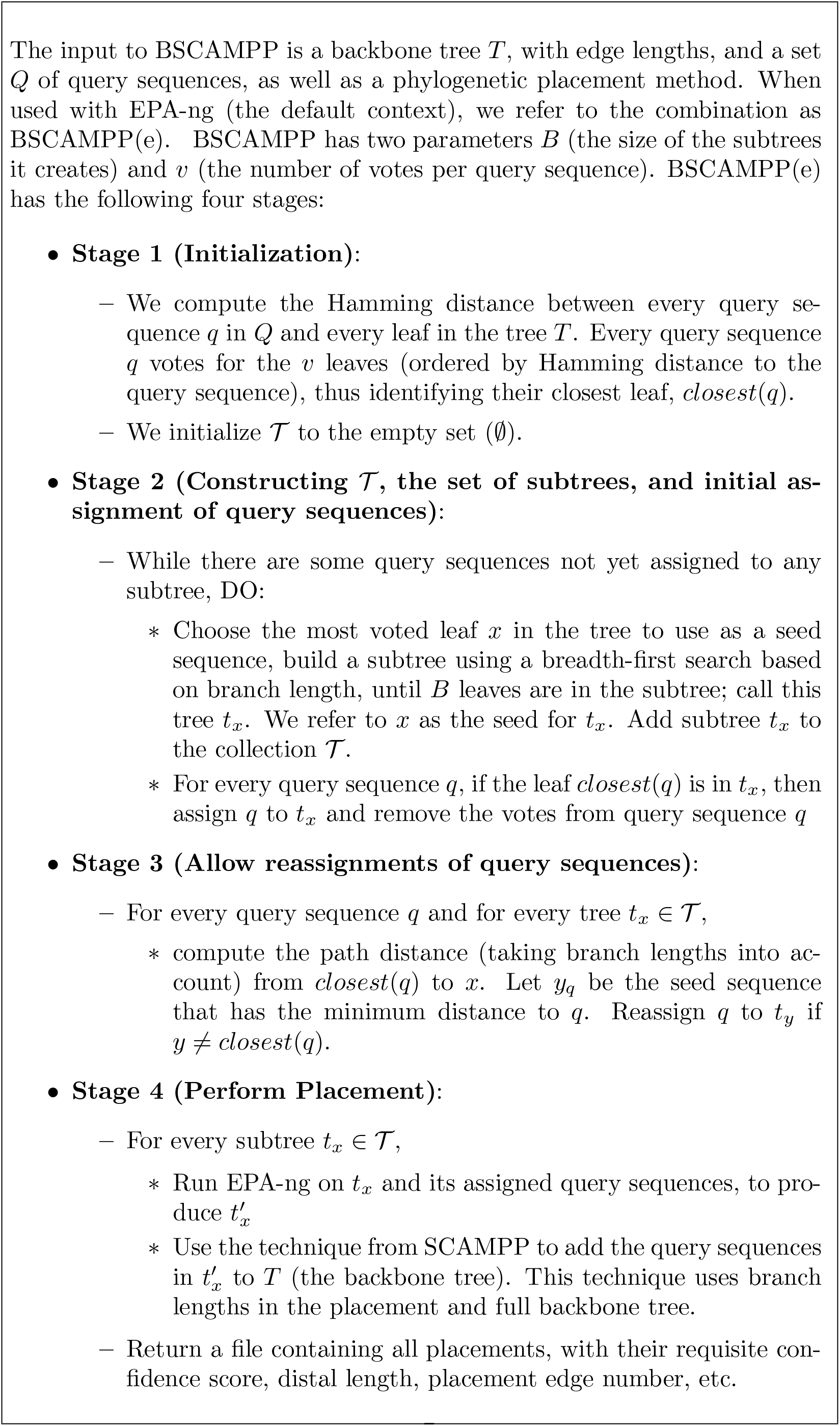

### Implementation details

On top of EPA-ng’s parallel implementation, BSCAMPP(e) is written in Python with certain parts written using OpenMP in C++. Since the Hamming distance computation is a computationally intensive portion of the BSCAMPP(e) framework (requiring 𝒪 (*rql*) for *q* queries of length *l* compared to *r* reference leaves), a parallel implementation using OpenMP allows for easier batch processing of queries.

## 4 Performance Study Design

### 4.1 Overview

We evaluate placement methods for use with short sequences, the application similar to what would be encountered in placing short reads into phylogenetic trees or taxonomies. Since many reads are characterized in each run, scalability to large numbers of reads is a relevant question. We use both simulated and biological datasets in this study, dividing the datasets into training data and testing data.

Our training data are simulated datasets with known true trees, and the testing data are both biological datasets (with estimated maximum likelihood trees) as well as simulated datasets. We report placement error using “delta error”, a standard metric used in prior studies [11, 18] (see Section 4.7).

In addition to the preliminary experiments (i.e., Experiment 0, reported in Section 2), we ran the following experiments for this study:

- Experiment 1 is the design of BSCAMPP(e) (i.e., BSCAMPP used with EPA-ng) on training data.
- Experiment 2 compares BSCAMPP(e) to other phylogenetic placement methods on testing data using estimated alignments.
- Experiment 3 compares BSCAMPP(e) to other phylogenetic placement methods using reads with sequencing error.
- Experiment 4 evaluates BSCAMPP(e) on datasets with changing rates of evolution.
- Experiment 5 evaluates the scalability of BSCAMPP(e) on a large backbone tree (180,000 leaves) from the test data and varying the number of query sequences.

See Supplementary Materials S4 for additional details about datasets and commands used in our experimental study.

### 4.2 Methods

In Experiments 1–5, we compare EPA-ng (v0.3.8), UShER, App-SpaM, and RAPPAS, APPLES-2 (v2.0.11), SCAMPP (v2.1.1) used with pplacer or EPA-ng (i.e., SCAMPP(p) and SCAMPP(e)), and BSCAMPP used with EPA-ng (i.e., BSCAMPP(e)). Within SCAMPP(p) we use pplacer (v1.1.alpha19) with taxtastic [19] for placement, since this improves accuracy and scalability according to [3, 12].

Some phylogenetic placement methods require numeric parameters to be estimated for the backbone trees. All backbone tree numeric parameters (branch lengths, 4 × 4 substitution rate matrix, etc.) are re-estimated according to the specifications of the phylogenetic placement method: RAxML-ng (v1.0.3) [20] for EPA-ng, UShER, App-SpaM, SCAMPP(e), and BSCAMPP(e), and FastTree-2 (v2.1.11) [17] under Minimum Evolution for APPLES-2.

### 4.3 Computational resources

All methods are given four hours to run with 64 GB of memory and 16 cores. These analyses were run on the UIUC Campus Cluster, which is heterogeneous (i.e., some machines are older and hence slower than others). While all methods are given 16 cores and 64 GB, any given analysis may have access to very different computational resources. For SCAMPP(e), SCAMPP(p), and UShER, when placement time was over four hours (which occurred in all experiments with 10,000 query sequences), the query sequences were split into 40 subsets of 250 sequences each. SCAMPP(e), SCAMPP(p), and UShER were then run 40 times for each subset containing 250 query sequences.

### 4.4 Datasets

All datasets are from prior studies and are freely available in public repositories.

#### 4.4.1 RNASim

We use samples from the RNASim dataset [16], which is a simulated dataset of 1,000,000 sequences that evolve down a model tree under a biophysical model to preserve RNA secondary structure. Subsets of the million-sequence simulated dataset were used in prior studies evaluating phylogenetic placement methods [10–12], and provide a substantial challenge due to the dataset size. For this study, we split this dataset into two subsets by taking the model tree and splitting it into two clades, with one having approximately 600,000 sequences and the other having approximately 400,000 sequences. This defines two sets of sequences, with the smaller one used for training (Experiment 1) and the larger one for testing (Experiments 2, 3, and 5). We place into a maximum likelihood tree on the true alignment (estimated using FastTree-2), using the true alignment for our Experiments 1 and 5 and using alignments estimated with UPP [21] for Experiments 2 and 3.

#### 4.4.2 nt78

We also use the nt78 dataset, which were simulated for FastTree-2 [17]; these contain 10 replicates, simulated with Rose [22], each with 78,132 sequences in a multiple sequence alignment and the simulated backbone tree. We picked one replicate randomly, using 68,132 sequences for the backbone and 10,000 sequences for the query sequences. We placed the query sequences into a maximum likelihood tree, estimated using FastTree-2 [17] on the true alignment. This dataset is used in the training phase (Experiment 1).

#### 4.4.3 16S.B.ALL

For biological dataset analysis, we use 16S.B.ALL, a large biological dataset with 27,643 sequences and a reference alignment based on structure from The Comparative Ribosomal Website (CRW) [23]. 16S.B.ALL contains some duplicate sequences; these were removed before analysis, producing a dataset with 25,093 sequences. Of these, 5,093 sequences were randomly selected as query sequences and the remaining were made backbone sequences. A maximum likelihood tree for this dataset was computed for the SATé-II [24] study on this reference alignment using RAxML [25] and serves as the backbone tree into which we place the query sequences. When computing delta error, we used the 75% bootstrap tree (i.e., the result of collapsing all edges with support below 75%) as the reference topology. The maximum likelihood tree and the 75% bootstrap tree are available at [26].

#### 4.4.4 5000M(2-4)

This dataset, originally from [27] was generated using INDELible with a heterogeneous indel model. Each set contains 5000 simulated sequences. The 5000M2-het condition reflects the highest rate of evolution in the dataset, and the 5000M4-het condition reflects the lowest rate of evolution. For our experiments, 1000 sequences are randomly selected as queries, and used to generate 10,000 fragmentary sequences. We place these queries into a maximum likelihood tree, estimated using FastTree-2 [17] on the true alignment of the remaining 4000 sequences.

### 4.5 Fragmentary sequence generation

For Experiments 1, 2 and 5 we generated fragments from the full-length sequences starting at a randomly selected location. The fragmentary sequences are a mean of 10% of the original ungapped sequence length with a standard deviation of 10 nucleotides.

For Experiments 3 and 4 we simulate reads with sequencing error. Illumina length reads are generated using the ART sequence simulator [28]. We also generated PacBio length reads with higher sequencing error using PBSIM [29].

### 4.6 Additional details about experiments

In Experiment 1, we use ∼400, 000 sequences from the RNASim dataset, from which we randomly select 50,000 sequences to define the backbone sequences, 10,000 sequences for query sequences, and the remaining 340,000 sequences are not used. Sites of the true alignment on the 60,000 sequences containing more than 95% gaps are removed. To define the backbone tree on the set of sequences, we run FastTree-2 under the GTRGAMMA model on the true alignment.

For Experiments 1 and 5 the true alignments of our fragmentary sequences are used. In testing experiments 3 and 4, reads with sequencing error are used. For Experiments 2–5 we align all reads to the true backbone alignment using UPP [21]. See supplement for command settings and details.

For the nt78 datasets, we use the third replicate for the experiment. As the true alignment is not very gappy, we do not perform any masking. We pick 10,000 sequences at random for the query sequences and use the remaining 68,132 sequences as backbone sequences. We also define the backbone tree by running FastTree-2 on the true alignment of the backbone sequences.

### 4.7 Evaluation Criteria

We report placement error using average delta error [10, 12, 18], where the delta error for a single query sequence is the increase in the number of missing branches (FN) produced by adding the query sequence into the backbone tree, and hence is always non-negative; this is the same as the node distance when the backbone tree is the true tree. This requires the definition of the “true tree”, which is the model tree for the simulated data and the published reference tree for the biological data. The methods are also evaluated with respect to runtime and peak memory usage.

## 5 Results

### 5.1 Experiment 1: BSCAMPP(e) design

In designing BSCAMPP, we considered four different strategies, described in the Supplementary Materials Section S2.1. We used the training data for this exploration. In comparing the variants for speed and accuracy, we found that variant 4 (see Section 3) provided accuracy that was comparable with the next most accurate method, but had better computational performance (Figures S3 and S4 in the Supplementary Materials). Based on this, we selected variant 4. Having selected variant 4, we then performed additional experiments on the two training datasets to set the values for two parameters: the size of the subtrees and the number of votes per query sequence. We varied the subtree parameter setting from 1000, 2000, 3000, 5000 and 10,000 leaves. For each subtree size, we ran BSCAMPP(e) with 5 and 25 votes per query sequence. Results for this experiment are shown in Figure 1 (see also the Supplementary Materials Tables S1 and S2).

**Figure 1:**
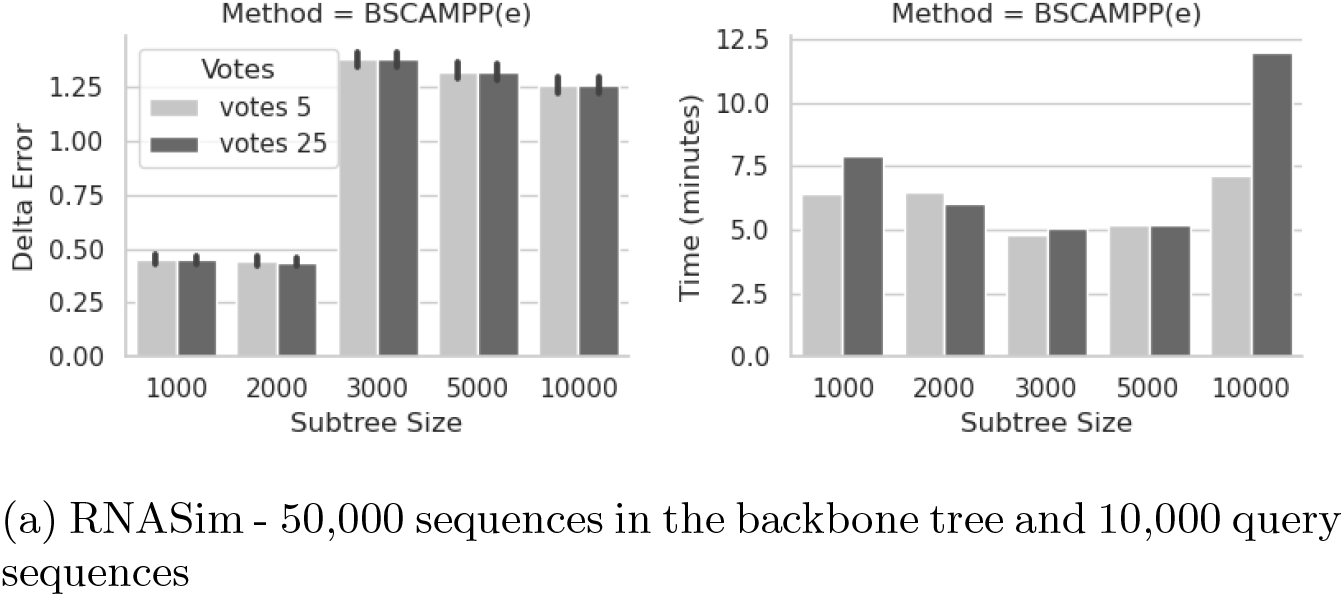
Experiment 1 - Subtree Size and Vote settings for BSCAMPP(e) on training dataset using the true alignment of the queries. Mean Delta Error plus standard error (left) and total runtime (right) for placement of all 10,000 fragmentary query sequences. We show placement time and error for BSCAMPP(e) varying parameter *v* (the number of votes per query) and the parameter *B* (the size of the subtree). The fragmentary sequences are a mean of 10% of the original ungapped sequence length with a standard deviation of 10 nucleotides.

**Figure 2:**
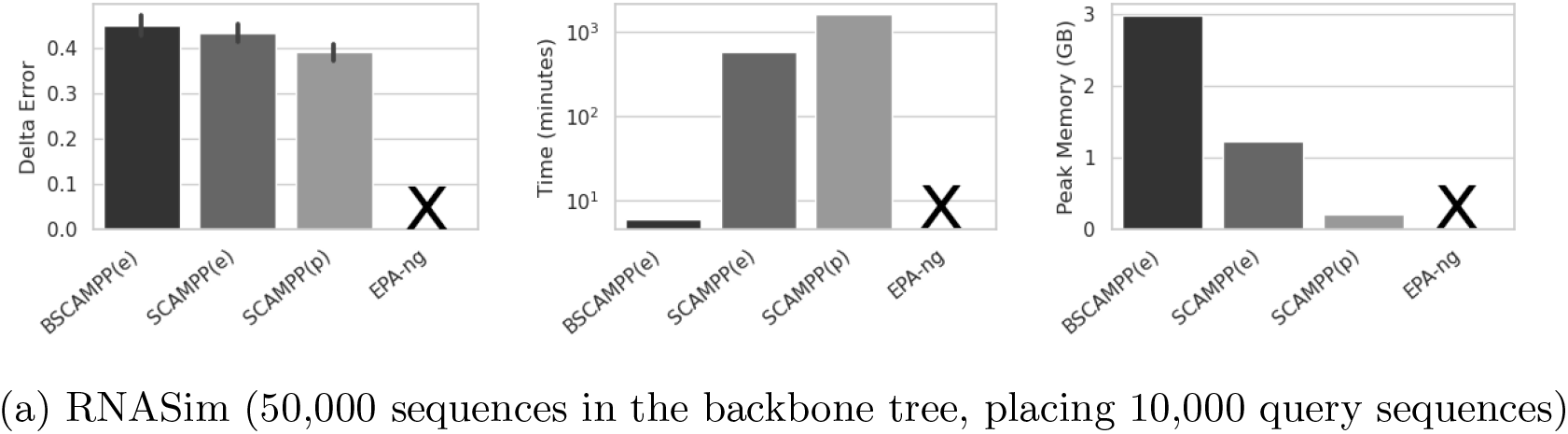
Experiment 1: Method Comparison on the training dataset on true query sequence alignments. Mean delta error (left), runtime (center), and peak memory usage in GB (right) for placement of all 10,000 query sequences on the RNASim backbone trees with 50,000 leaves. We show placement time for BSCAMPP(e) for 5 votes using a subtree of 2000 leaves. SCAMPP(e) and SCAMPP(p) similarly use a subtree size of 2000 leaves. The results from SCAMPP are also included for runtime, peak memory usage, and delta error. EPA-ng was not shown because it was unable to run given 64 GB of memory and 16 cores on these datasets due to out-of-memory issues.

Note that when the subtree size increases over 2,000 leaves, BSCAMPP(e) has over twice the error than it does for 1,000- and 2,000-leaf subtrees. The number of votes does not produce a large change in delta error. The runtime shows that the placements with all of the subtree sizes and votes were able to complete in under 15 minutes. For both subtree sizes where the delta error is low, using 5 votes for parameter *v* results in faster or similar runtimes.

Based on these trends a subtree size of 2,000 and 5 votes are chosen as defaults for parameters *B* and *v*, respectively.

Using these parameter settings for BSCAMPP(e) we show a comparison to both SCAMPP variants (SCAMPP(e) used with EPA-ng and SCAMPP(p) used with pplacer) and EPA-ng in Figure 2. EPA-ng was unable to place into our 50,000 leaf backbone tree given the memory limitations. SCAMPP(p) is the most accurate method shown with a delta error of 0.39, followed by SCAMPP(e) at 0.43, and BSCAMPP(e) at 0.45. While there is an improvement in accuracy with SCAMPP(p) it requires a combined effort of over 27 hours to place all sequences, whereas BSCAMPP(e) used only 6 minutes.

### 5.2 Experiment 2: Method comparison using estimated alignments

We selected App-SpaM and RAPPAS, the alignment-free methods studied in [14] that had the best accuracy, to evaluate suitability for placing into large trees, and compared them to alignment-based methods UShER, APPLES-2, BSCAMPP(e), and EPA-ng. This experiment was performed on our testing datasets: nt78, RNASim 50K, and 16S.B.ALL. All query sequences are fragmentary with length 10% of the full length. As with all our experiments, the methods were given 64GB of memory.

The results when query sequences are placed with an alignment using UPP [21] (rather than with the true alignment as in Experiment 1) are shown in Figure 3 (see Figures S1 and S2 for results given true alignments of query sequences). These results showed that RAPPAS and EPA-ng failed to complete on these datasets (except for EPA-ng on 16S.B.ALL) due to high memory requirements.

**Figure 3:**
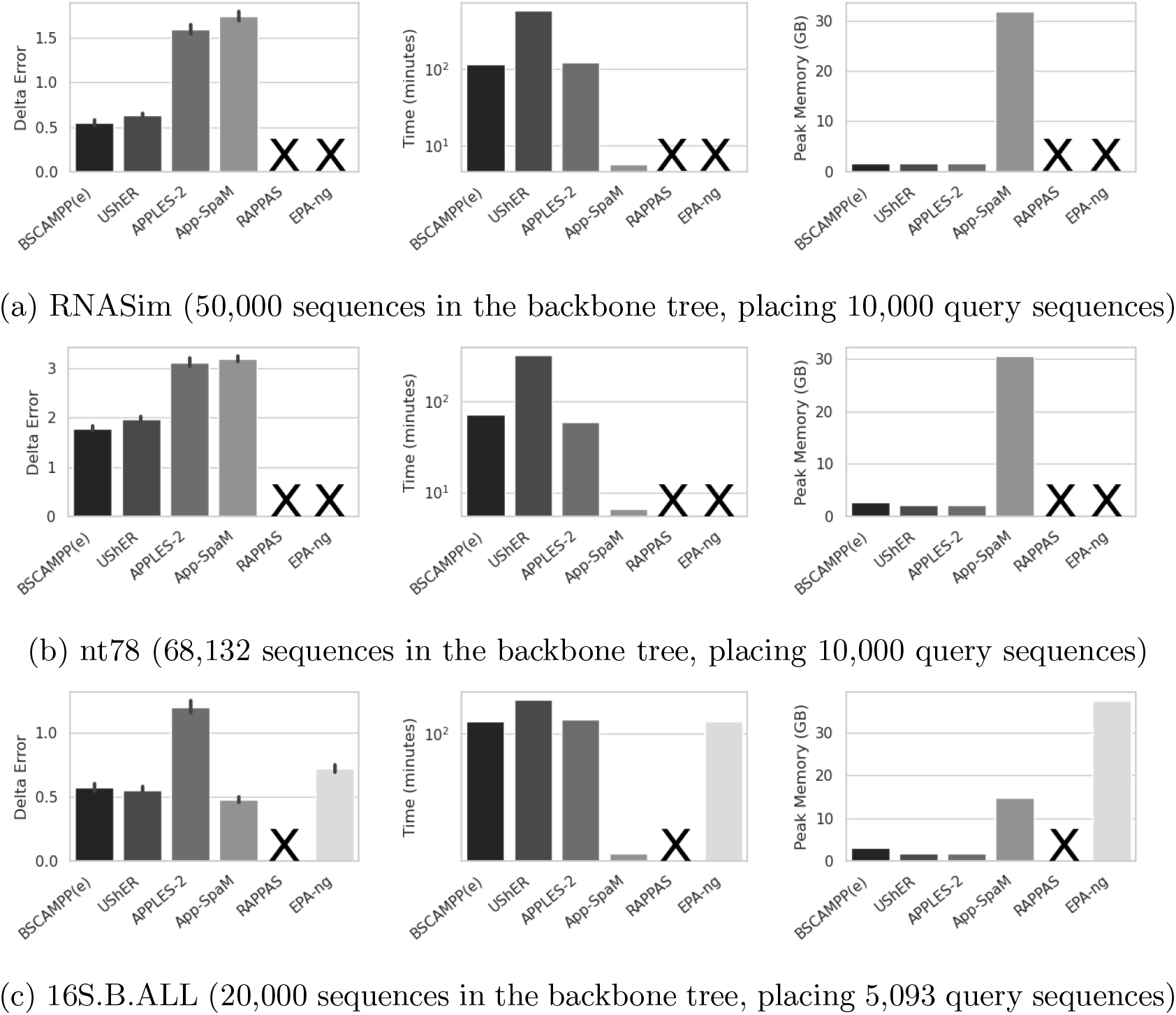
Experiment 2: Method Comparison using estimated query sequence alignments on testing data. We show from left to right - Mean delta error, total runtime, and peak memory usage in GB for three datasets placing large sets of queries into large reference trees. Fragmentary query sequence alignments are estimated using UPP, and the alignment time is included in the runtime for the alignment-based methods (all except RAPPAS and App-SpaM). We show placement time for BSCAMPP(e) for 5 votes with a subtree size of 2000 (default settings). The results from UShER, APPLES-2, App-SpaM, RAPPAS, and EPA-ng are also included (an **X** indicates that RAPPAS and EPA-ng were unable to run due to out-of-memory issues).

App-SpaM was able to complete on the nt78 and RNASim 50K datasets, but had similar accuracy to APPLES-2 on both datasets (in turn, APPLES-2 was much less accurate than BSCAMPP(e), and UShER). App-SpaM was the fastest method, due to the other methods requiring alignments of the query sequences, and the vast majority of the runtime for APPLES-2 and BSCAMPP(e) is the calculation of the alignment of the query sequences to the backbone alignment (see Tables S1 and S2). However, App-SpaM was unable to complete on the RNASim 200K dataset, and had higher memory requirements than the other tested methods on these datasets.

### 5.3 Experiment 3: Method comparison using reads with sequencing error

Method evaluation is shown on both Illumina and Pacbio style simulated reads for three of our testing datasets in Experiment 3. We show results for BSCAMPP(e) compared to two other alignment-based methods, APPLES-2 and UShER, and the alignment-free method, App-SpaM, which performed best in Experiment 2.

Figure 4, shows a few trends for placement of Illumina-style reads. For both simulated datasets where ground truth is known, BSCAMPP(e) is the most accurate method, followed by UShER, APPLES-2, and then App-SpaM. On the biological dataset (16S.B.ALL), App-SpaM is the most accurate method, and BSCAMPP(e) and UShER closely follow. APPLES-2 shows over twice the placement error of BSCAMPP(e) and UShER.

**Figure 4:**
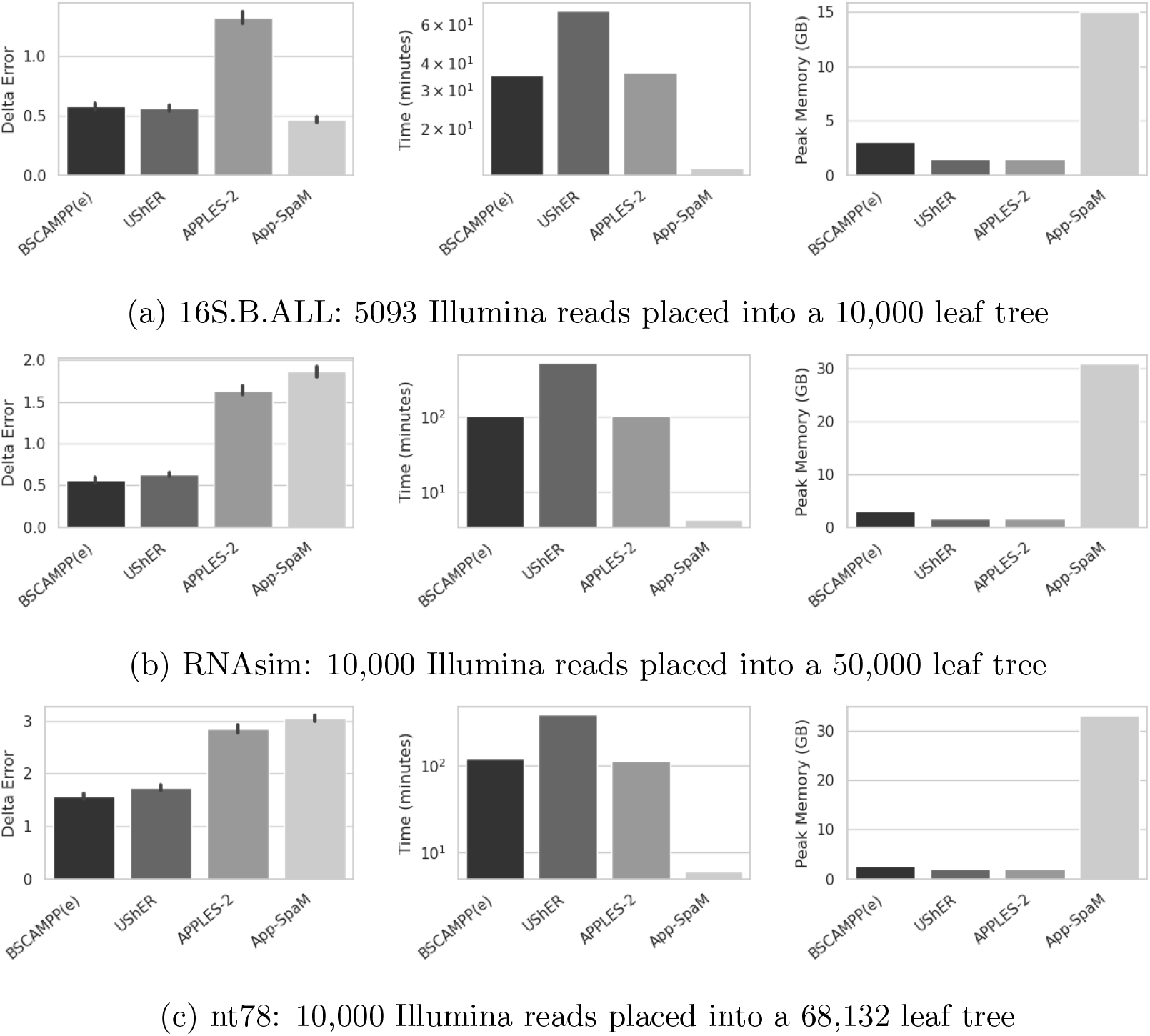
Experiment 3: Mean delta error, total runtime, and peak memory usage for phylogenetic placement methods (BSCAMPP(e), APPLES-2, UShER, and App-SpaM) on three testing datasets using **Illumina** style sequencing reads with sequencing error. For BSCAMPP(e), APPLES-2, and UShER the reads are first aligned to the reference alignment using UPP, and the aligned queries are placed into the reference alignment and tree.

Runtime for all alignment-based methods includes the alignment process. As in Experiment 2 this accounts for the majority of the runtime for both BSCAMPP(e) and APPLES-2. UShER is the slowest method shown and App-SpaM is the fastest, with near instantaneous placement. The peak memory usage for App-SpaM shows that it uses more memory than the alignment-based methods, requiring 30 GB for both simulated datasets.

The Pacbio-style reads show similar trends. Here in all cases BSCAMPP(e) is the most accurate method, closely followed by UShER. APPLES-2 has more than twice the error of BSCAMPP(e) and UShER, and App-SpaM has the highest error shown. Again similar runtime trends to the Illumina-style reads hold. App-SpaM places all queries in minutes, while the runtime of the alignment process increases the runtime for all other methods shown. BSCAMPP(e) and APPLES-2 both have runtime that is dominated by the alignment method and UShER is the slowest method. Here BSCAMPP is showing slightly more memory usage, than UShER and APPLES-2 at up to 4 GB for the biological dataset. App-SpaM still uses the most memory of all methods shown.

### 5.4 Experiment 4: Method comparison on datasets with variable rates of evolution

The purpose of Experiment 4 is to evaluate BSCAMPP(e) in comparison to other placement methods on the testing datasets including a a dataset with varying rates of evolution. Experiments 2 and 3 showed that both BSCAMPP and UShER have similar accuracy, and are the most accurate methods tested. UShER was designed first for use with large COVID phylogenies, and thus has been tested in conditions with high sequence similarity. In Experiment 4, we evaluate BSCAMPP, UShER, APPLES-2, and App-SpaM using datasets with varying rates of evolution to further understand the difference between the performance.

Figure 6 shows that for all conditions, 5000M2 (highest rate of evolution) through 5000M4 (lowest rate of evolution), and for both read lengths, BSCAMPP(e) is the most accurate method. The Illumina-style reads show the largest range in accuracy between methods, on the 5000M2 data with the highest rate of evolution. Here BSCAMPP(e) at an error of *<* 1 has less than half the error of UShER at around 2, the second most accurate method. APPLES-2 shows an increase in error to almost 10, and finally App-SpaM’s error is over 15. As the rate of evolution lowers, these trends continue but all methods improve in accuracy. Pacbio reads show a less clear delineation between the accuracy of BSCAMPP(e) and UShER, but BSCAMPP(e) is still more accurate, closely followed by UShER and APPLES-2. App-SpaM shows higher error than all other methods.

**Figure 5:**
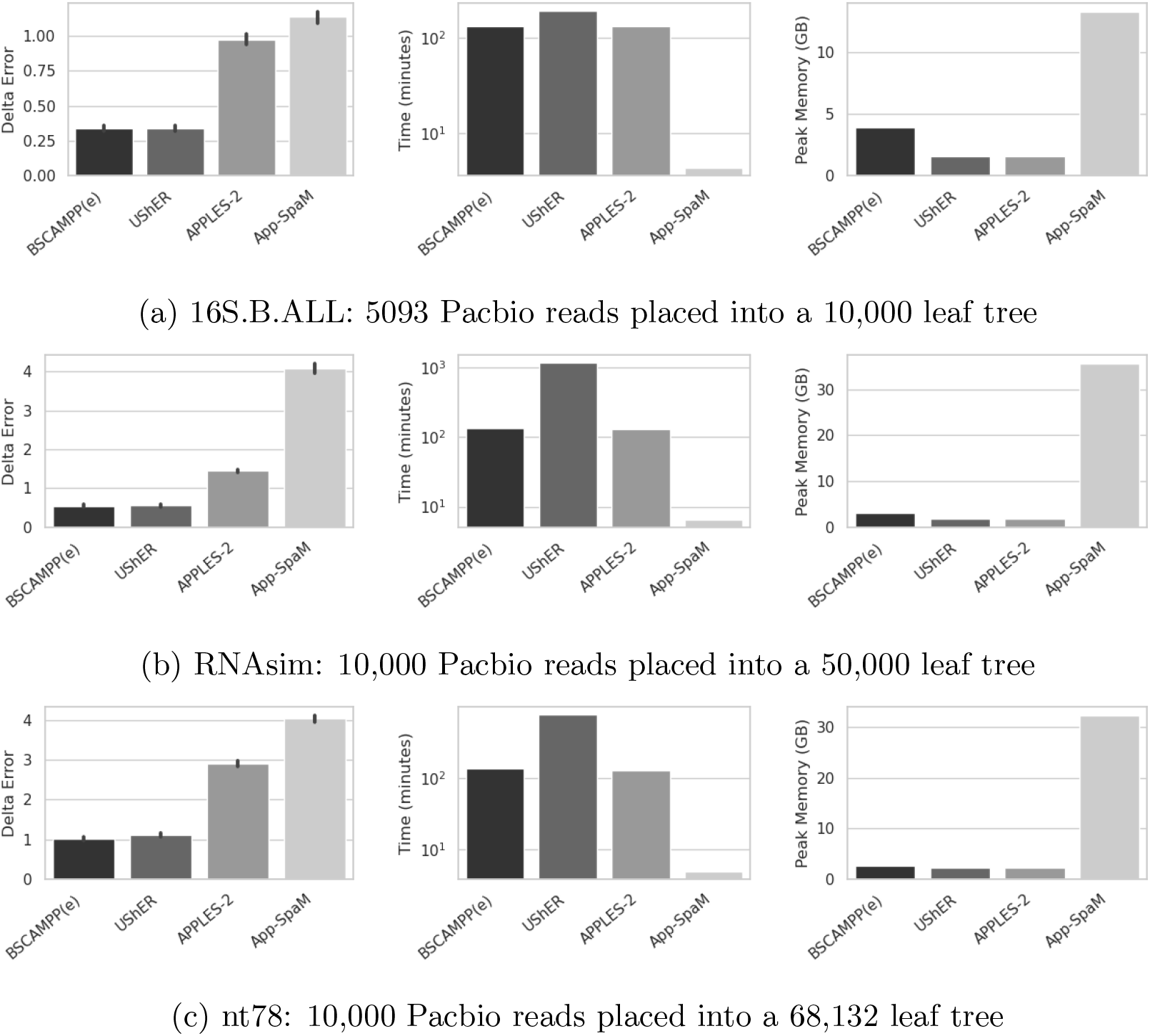
Experiment 3: Mean delta error, total runtime, and peak memory usage for phylogenetic placement methods (BSCAMPP(e), APPLES-2, UShER, and App-SpaM) on three testing datasets using **Pacbio** style sequencing reads with sequencing error. For BSCAMPP(e), APPLES-2, and UShER the reads are first aligned to the reference alignment using UPP, and the aligned queries are placed into the reference alignment and tree.

**Figure 6:**
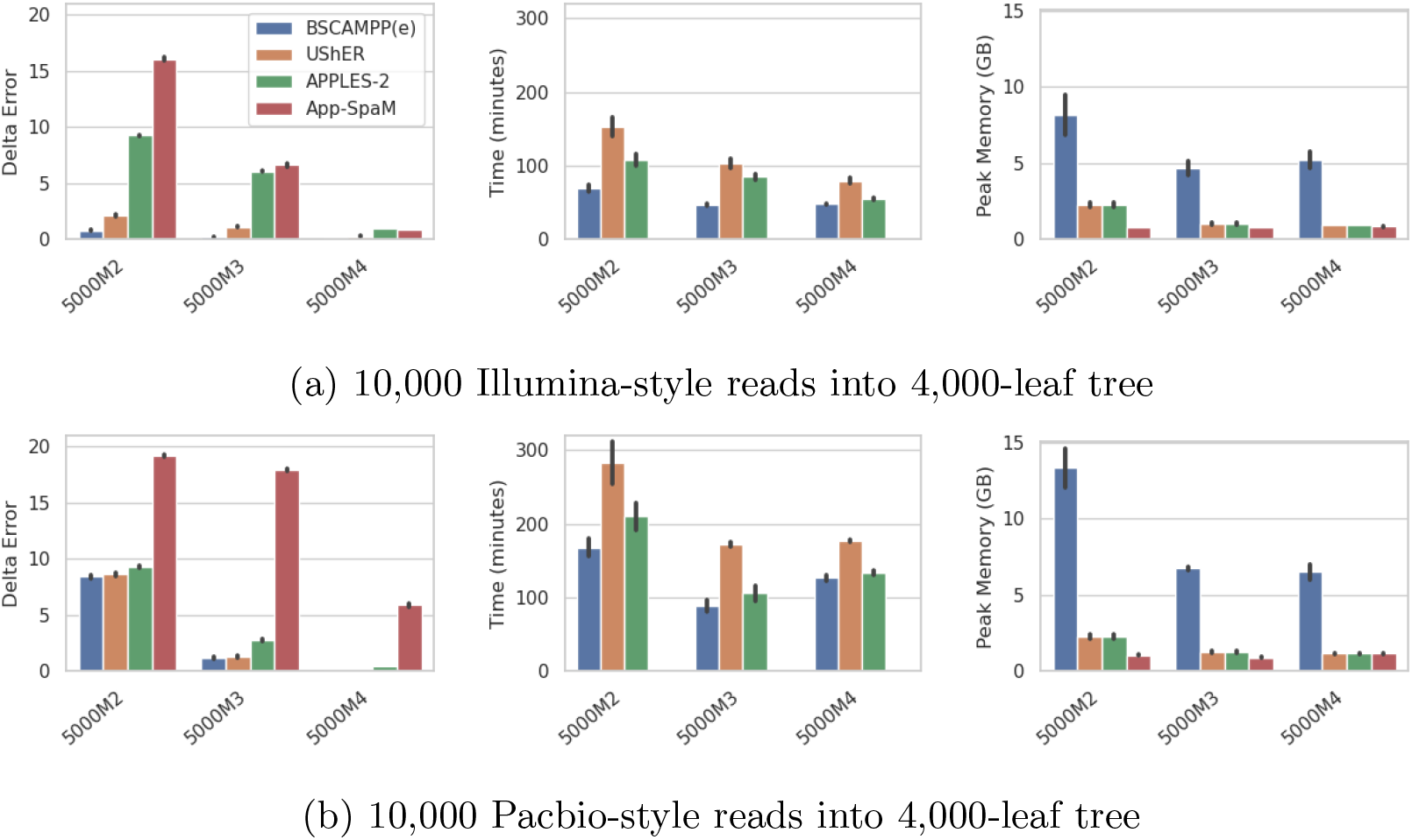
Experiment 4: 5000M2-4: Three simulated datasets with varying rates of evolution from higher (5000M2) to lower (5000M4). Shows placement of 10,000 queries into 4,000-leaf trees on two replicates for each dataset. Delta error is shown only for sequences that all methods successfully placed. From left to right: Mean delta error, total runtime, and peak memory usage for phylogenetic placement methods (BSCAMPP(e), UShER, APPLES-2, and App-SpaM) on three testing datasets using **Illumina** style sequencing reads with sequencing error (above) and **Pacbio** style sequencing error (below). For BSCAMPP(e), APPLES-2, and UShER the reads are first aligned to the reference alignment using UPP, and the aligned queries are placed into the reference alignment and tree.

The alignment-based methods have the alignment phase included in the runtime. Of the alignment-based methods BSCAMPP(e) is the fastest shown, followed by APPLES-2 and then UShER. App-SpaM is near instantaneous (and not even visible in Figure 6), and here requires the least memory usage.

BSCAMPP(e) uses more memory here than all other methods at up to 15 GB.

We note that APPLES-2 did not return placements for all queries for many of these datasets. We show delta error on placements for all query sequences that returned placements for all methods here. For a reporting on the number of query sequences APPLES-2 did not generate placements for please see Supplementary Materials Table S3. For results on all 10,000 queries (without APPLES-2) see Supplementary Figure S5.

### 5.5 Experiment 5: Method comparison on ultra-large trees

Our final experiment is designed to verify that BSCAMPP(e) scales to both large query sets and ultra-large backbone tree sizes. Additionally, we check that BSCAMPP(e) is able to achieve sub-linear runtime scaling with respect to the number of query sequences. We use RNASim 200K for this study, with a backbone tree having 180,000 leaves, and placing from 20 to 20,000 fragmentary query sequences into the tree. We limited this study to BSCAMPP(e) and APPLES-2, as these were the only methods that are able to place into the 50,000-leaf tree within 1 hour. App-SpaM failed to complete on this dataset due to out-of-memory issues and is not included here.

Results in Figure 7 show that BSCAMPP(e) is able to place 20,000 sequences in just 25 minutes, whereas 20 sequences require 1 minute. The runtime for BSCAMPP(e) increases by a factor of 6.7, 2.5, and 1.5 each time the number of queries increases tenfold. Similarly, APPLES-2 shows a sub-linear curve; the runtime increases by a factor of 3.6, 3.8, and then 6.9 each time the number of queries increases tenfold. APPLES-2 is faster for 20, 200, and 2,000 query sequences, but BSCAMPP(e) is faster at 20,000 query sequences. Moreover, both methods are able to complete in under 1 hour, even when placing 20,000 query sequences into this large tree with 180,000 sequences. Finally, both methods use under 3 GB, well under the computational limit of 64 GB.

**Figure 7:**
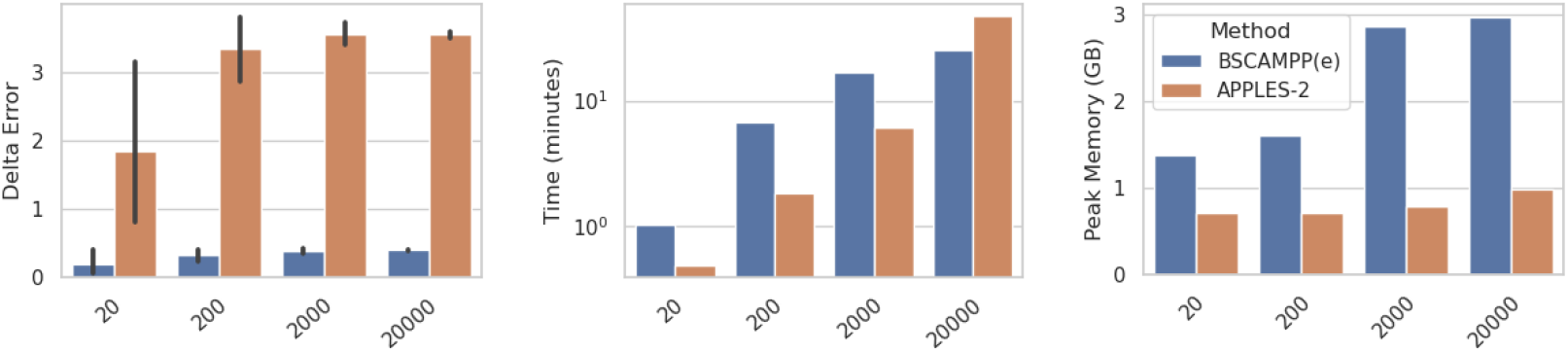
Experiment 5: RNASim 180k. Comparison of BSCAMPP(e) and APPLES-2 given ultra-large backbone tree (180,000 sequences) and true query sequence alignments. We show mean delta error (left), total runtime (center) for placement of all query sequences, and peak memory usage (right). We run each method for 20, 200, 2,000, and 20,000 query sequences. Only APPLES-2 and BSCAMPP(e) are shown, since they were the only methods able to complete in under four hours for 20,000 queries.

Results shown in Figure 7 also allow us to compare methods for accuracy, and see how changing the number of query sequences affects this. We see that BSCAMPP(e) delta error is not impacted by the number of query sequences, and when at least 200 sequences are considered the delta error for APPLES-2 is also not impacted; this makes sense. In addition, BSCAMPP(e) shows a much lower delta error than APPLES-2 (consistent with trends in previous experiments).

### 5.6 Computational performance

We focus now on runtime for only alignment-based methods, and note that the fastest method in all previous experiments has been the alignment-free method App-SpaM. This speed advantage for App-SpaM comes at an accuracy loss, so we seek to explore the runtime for only the placement phase of all other methods (without including the alignment phase).

When restricting our attention to APPLES-2 and BSCAMPP(e), we find that both were very fast, with no clear dominance of one over the other (Table 2). When considering only the time to do phylogenetic placement (and so assuming the query sequences are given in an alignment to the backbone sequences), APPLES-2 never used more than 47 minutes and BSCAMPP(e) never used more than 26 minutes.

UShER required more time than BSCAMPP and APPLES, but less than either of the SCAMPP methods, with the fastest time at 41 minutes on 16S.B.ALL (the dataset with the lowest average p-distance - see Table 1). We see that although SCAMPP(e), and SCAMPP(p) were able to run on all the datasets, they were by far the most computationally intensive of the tested methods. Furthermore, SCAMPP(e) used from 30 to 70 times as much time as BSCAMPP(e).

**Table 1:** Dataset Statistics— The first column gives the name of the dataset and the publication describing the dataset. For each dataset, we show the number of sequences, the length of the reference alignment, its type (biological or simulated), the mean and maximum p-distance (i.e., normalized Hamming distances) between pairs of sequences in the alignment, and the proportion of the alignment that is gapped.

| Dataset | number of sequences | alignment length | type | p-dist mean | p-dist max | gaps prop. |
| --- | --- | --- | --- | --- | --- | --- |
| 16S.B.ALL [23] | 27,643 | 6857 | bio | .210 | .769 | .118 |
| nt78 [17] | 78,132 | 1287 | sim | .404 | .639 | .006 |
| RNASim [16] | 1,000,000 | 1620 | sim | .410 | .618 | .051 |
| 5000M2 [27] | 5,000 | 52,606 | sim | .693 | .801 | .981 |
| 5000M3 [27] | 5,000 | 24,062 | sim | .660 | .765 | .958 |
| 5000M4 [27] | 5,000 | 22,403 | sim | .530 | .673 | .957 |

**Table 2:** Runtime results for placement phase across all datasets and backbone tree sizes for all alignment-based methods. These fragmentary queries were generated without sequencing error and the true alignment to the reference is used. An **X** indicates that the analysis failed due to out-of-memory issues, and a “-” indicates the analysis was not attempted due to the expectation of excessive runtime (exceeding 500 minutes) or excessive memory requirements (exceeding 64 GB).

| Method | Total Runtime (for all queries in minutes) |  |  |  |  |
| --- | --- | --- | --- | --- | --- |
|  | Training | Testing |  |  |  |
| Dataset: | RNASim | nt78 | RNASim |  | 16S |
| Backbone Size: | 50,000 | 68,132 | 50,000 | 180,000 | 20,000 |
| Nbr Queries: | 10,000 | 10,000 | 10,000 | 20,000 | 5,093 |
| BSCAMPP(e) | 6 | 14 | 7 | 26 | 4 |
| SCAMPP(e) | 580 | 429 | 486 | - | 215 |
| SCAMPP(p) | 1636 | 1426 | 1421 | - | 952 |
| EPA-ng | X | X | X | - | 17 |
| USHER | 510 | 276 | 444 | - | 41 |
| APPLES-2 | 6 | 4 | 5 | 47 | 9 |

We also note that BSCAMPP(e) is able to run on datasets where EPA-ng fails due to out-of-memory issues. Indeed, the only large dataset in our study that EPA-ng was able to complete was the 16S.B.ALL dataset, and on this dataset, EPA-ng has a higher error than BSCAMPP(e).

The memory usage was never that high for any method that succeeded in running (the most was EPA-ng, at 37 GB), but as no method had access to more than 64 GB, it is possible that EPA-ng might have succeeded in running if it had access to a larger amount of memory.

## 6 Discussion

Our study established that RAPPAS, App-SpaM, and EPA-ng had high memory requirements, making all of these methods unable to scale to the largest dataset we examined (RNASim 200K, with 180K sequences in the backbone tree and 20,000 query sequences). Indeed, EPA-ng and RAPPAS were unable to run on the RNASim 50K dataset, due to their memory requirements. Thus, these three methods had higher memory requirements than the remaining methods, which included APPLES-2, UShER, and the SCAMPP- or BSCAMPP-enabled methods. This finding is perhaps as expected, as all the SCAMPP-, BSCAMPP-, and APPLES-2 methods use divide-and-conquer, limiting the phylogenetic placement effort to small datasets (at most 2000 leaves in the relevant placement subtree), and UShER’s mutation annotated tree reduces the memory requirements.

APPLES-2, BSCAMPP(e), and App-SpaM were the fastest methods, with a definite advantage to App-SpaM when the time to compute the alignment of the query sequences to the backbone alignment is considered. However, as noted above, App-SpaM did not complete the largest dataset. SCAMPP(p), SCAMPP(e) and UShER were the slowest of the methods we tested, as these methods incrementally place into the reference tree.

Our experiments consistently showed that all likelihood-based and parsimony-based placement methods had the lowest delta error of the tested methods, with relatively minor differences between them (although EPA-ng was less accurate than the other likelihood-based methods in some cases), making computational performance the main distinguishing feature. The only exception to this was App-SpaM on the 16S.B.ALL dataset when there was low or no sequencing error and where the reference tree used for delta error computation was the 75% bootstrap tree (i.e. not fully resolved).

Given all this, the result that BSCAMPP(e) is much faster than the other likelihood-based methods on large datasets is noteworthy. We also observed that BSCAMPP(e) improves the accuracy of APPLES-2, in many cases dramatically, and even improves on speed when the number of query sequences is large enough (at smaller numbers of query sequences, APPLES-2 is faster). However, if speed of placement is not of concern, then our experiments show that SCAMPP(p) is the most accurate method tested.

To understand why we see these trends, we consider what we learned about EPA-ng scalability on large backbone trees, both in terms of computational performance but also accuracy. Prior studies have suggested limits for EPA-ng to relatively small backbone trees due to computational reasons [10–12], but our Experiment 1 showed that EPA-ng had a jump in placement delta error as we increased the subtree size for the RNASim dataset. Thus, our study suggests potentially that EPA-ng may have some numeric issues when placing into very large trees that result in increased placement error, a trend that has been previously observed for pplacer [10]. Thus, we would hypothesize that when placing into very large trees, even if EPA-ng manages to finish it may have reduced accuracy due to numeric issues, and that BSCAMPP(e) can have higher accuracy than EPA-ng since it restricts the placement subtrees for EPA-ng to be relatively small (at most 2000 sequences).

From a computational viewpoint, the main issue for EPA-ng is its large memory usage, which can explain why EPA-ng is limited to backbone trees of at most moderate size. Thus, by limiting the placement tree size to at most 2000 sequences, not only does BSCAMPP(e) succeed in maintaining good accuracy, but it also ensures good scalability to very large backbone trees.

Our study may suggest that BSCAMPP(e) could be able to scale to much larger numbers of query sequences while maintaining the sublinear runtime. The basis for this optimism is that EPA-ng has been shown to place millions of queries [5] into smaller trees of a few hundred sequences, yet we have limited this study to larger backbone trees with at most 20,000 query sequences. The other point is that Experiment 5 suggests that the full efficiency of BSCAMPP(e) has not yet been reached at 20,000 query sequences, and that it may continue to be able to place even larger numbers of query sequences with runtimes that grow sublinearly with that number even for backbone trees of this size.

## 7 Conclusions

Phylogenetic placement into large backbone trees is fundamental to several bioinformatics problems, including microbiome analysis (e.g., taxonomic characterization of shotgun sequencing reads) and updating large phylogenetic trees. Yet, only a very small number of methods provide acceptable accuracy and scalability. SCAMPP [10] was designed to improve scalability of likelihood-based phylogenetic placement methods to large backbone trees, and enabled good accuracy in this context. However, by design SCAMPP did not ensure that the subproblems resulted in many query sequences being placed into the same subtree, which meant that it could not take advantage of EPA-ng’s ability to place many query sequences with a runtime that is sublinear in the number of query sequences. Our redesign, which we call BATCH-SCAMPP (or BSCAMPP for short) matches the accuracy of the SCAMPP and scalability to large backbone trees, and when used with EPA-ng (i.e., BSCAMPP(e)) achieves sublinear runtime in the number of query sequences. Moreover, BSCAMPP is extremely fast, nearly as fast as APPLES-2 for placing a moderate number of query sequences into a large backbone alignment, and slightly faster when placing large numbers of query sequences into very large backbone trees. Thus, BSCAMPP is the first likelihood-based phylogenetic placement method that is both highly accurate (due to its use of likelihood) and scalable in both dimensions – the number of query sequences and the size of the backbone tree (due to its use of divide-and-conquer and the reliance on EPA-ng for placement of many query sequences into small subtrees).

This study leaves several directions for future research. A more extensive study should explore phylogenetic placement of full-length sequences, and possibly also consider the problem of placing genome-length sequences. This particular direction raises issues of heterogeneity across the genome, a problem that is addressed by the DEPP [2] method for phylogenetic placement. In addition, while BSCAMPP is fast, a possible improvement for the runtime can be explored by implementing parallel processing of subtrees (i.e., running instances of EPA-ng in parallel for different query-subtree sets). This might be particularly helpful in cases with few queries per subtree. Future work should include an analysis of the runtime and memory usage impacts of running EPA-ng concurrently on multiple compute nodes, in addition to running multiple instances of EPA-ng using fewer threads on a single compute node.

There are applications of phylogenetic placement methods that could be improved through the use of larger backbone tree sizes and many query sequences, now enabled with better accuracy through BSCAMPP. One such application is that of *de novo* phylogenetic tree building. Studies have shown that scalable phylogeny estimation methods have suffered in the presence of sequence length heterogeneity, and phylogenetic placement may provide a more accurate alternative [30]. For example, the divide-and-conquer tree estimation pipeline GTM (Guide Tree Merger) [31] could benefit by using BSCAMPP. BSCAMPP can facilitate the initial tree decomposition of GTM for better placement of shorter, fragmentary sequences into an initial tree containing the longer full-length sequences, potentially leading to better final tree estimation.

Another application with very large backbone trees with many query sequences is taxon identification and abundance profiling. For example, TIPP [4] and its improved version TIPP2 [1] use phylogenetic placement (based on pplacer) to place query sequences into gene-specific taxonomies. As shown in [1], the accuracy improvement of TIPP2 over TIPP is mainly due to the increased size of the backbone trees in TIPP2 over TIPP, suggesting that the use of BSCAMPP(e) on large trees might lead to increased accuracy and reduced runtime compared to the current use of pplacer and smaller backbone trees.

## Data availability

All datasets used in this study are from prior publications.

## Acknowledgments

The authors acknowledge the financial support of the Department of Computer Science. EW was supported by a Siebel Scholars scholarship, a SURGE Fellowship, and a Wing Kai Cheng Fellowship. CS was supported by NSF grant 2006069 (to TW).

## Supplementary Materials

### S1 Additional Results for Experiment 0: Study with EPA-ng

**Figure S1:**
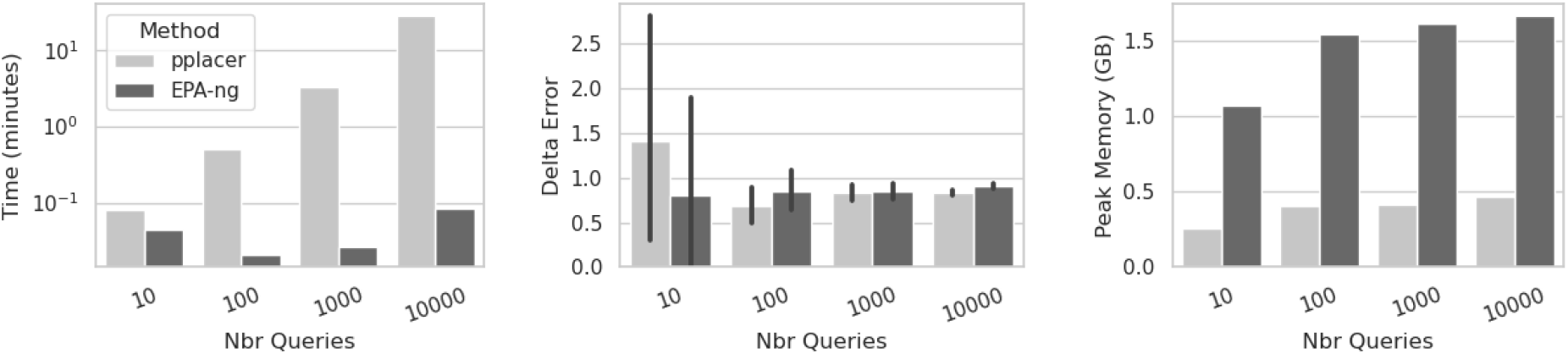
Experiment 0: Maximum-likelihood method comparison on training datasets on a 1000-leaf RNASim backbone tree. Runtime (left), Mean delta error (center), and peak memory usage in GB (right) are shown for placement of 10, 100, 1000, and 10,000 query sequences on an RNASim backbone tree with 1,000 leaves. Query sequences are fragmentary at 10% of the original sequence length. We show results for pplacer and EPA-ng (both methods that can be used within the SCAMPP and BSCAMPP frameworks).

The goal of this experiment was to understand the impact of the backbone tree size for EPA-ng using both fragmentary and full-length query sequences. We extracted a subset of the RNASim training dataset for this experiment. The subset is a clade with 10,200 leaves of the training RNASim dataset, from which we randomly selected 200 leaves as queries. For the remaining 10,000 leaves, we randomly selected from 1000 to 10,000 leaves to form the backbone tree. For the queries, we made them into fragments ranging from 10%-length to full-length. All queries were placed into the backbone trees using EPA-ng in one of two modes: “batch”, where we place all 200 queries all at once (a feature allowed by EPA-ng), and “one-by-one”, where we place each query independently by running EPA-ng on each query.

We show delta error on various backbone tree sizes, placement strategies (batch vs. one-by-one), and for either full-length sequences or fragmentary sequences (of lengths that are 10%, 25%, 50% of the full-length sequences) in Figure S2.

Note that placement accuracy is highest when placing full-length sequences, and that placing very short sequences (the standard situation in metagenomics) has the highest error rates. For all query sequence lengths, accuracy improves for one-by-one placement as the backbone size increases, indicating the beneficial impact of taxon sampling. Interestingly, this is not true for batch placement.

For batch placement, phylogenetic placement accuracy is very much impacted by query length, with full-length sequences placed well at all explored backbone tree sizes (up to 10,000-leaf trees). However, fragmentary sequences are only placed well on small backbone trees (up to 2,000 leaves). Interestingly, this is not the case when using one-by-one placement, where all phylogenetic placement error rates remained low, even for fragmentary sequences.

Since our interest is in placing short sequences (corresponding to read placement), we focus now on the results shown for queries of 10%-length. For these very short query sequences, we see a jump in delta error for “batch” from backbone tree size of 2000 to 5000 and up. This trend is also observed with less fragmentary queries (e.g., fragments between 25% and 50% of the original length), except that the gap in accuracy between the two strategies becomes smaller.

In sum, this experiment shows that EPA-ng performs very differently between the batch mode (the case where it is able to obtain a runtime advantage) and the one-by-one mode, and that it can be much more accurate in one-by-one mode when placing fragmentary sequences. The key lesson is that if we wish to obtain good accuracy in placing fragmentary sequences, we need to either use EPA-ng in one-by-one mode or keep the backbone tree at most 2000 sequences.

**Figure S2:**
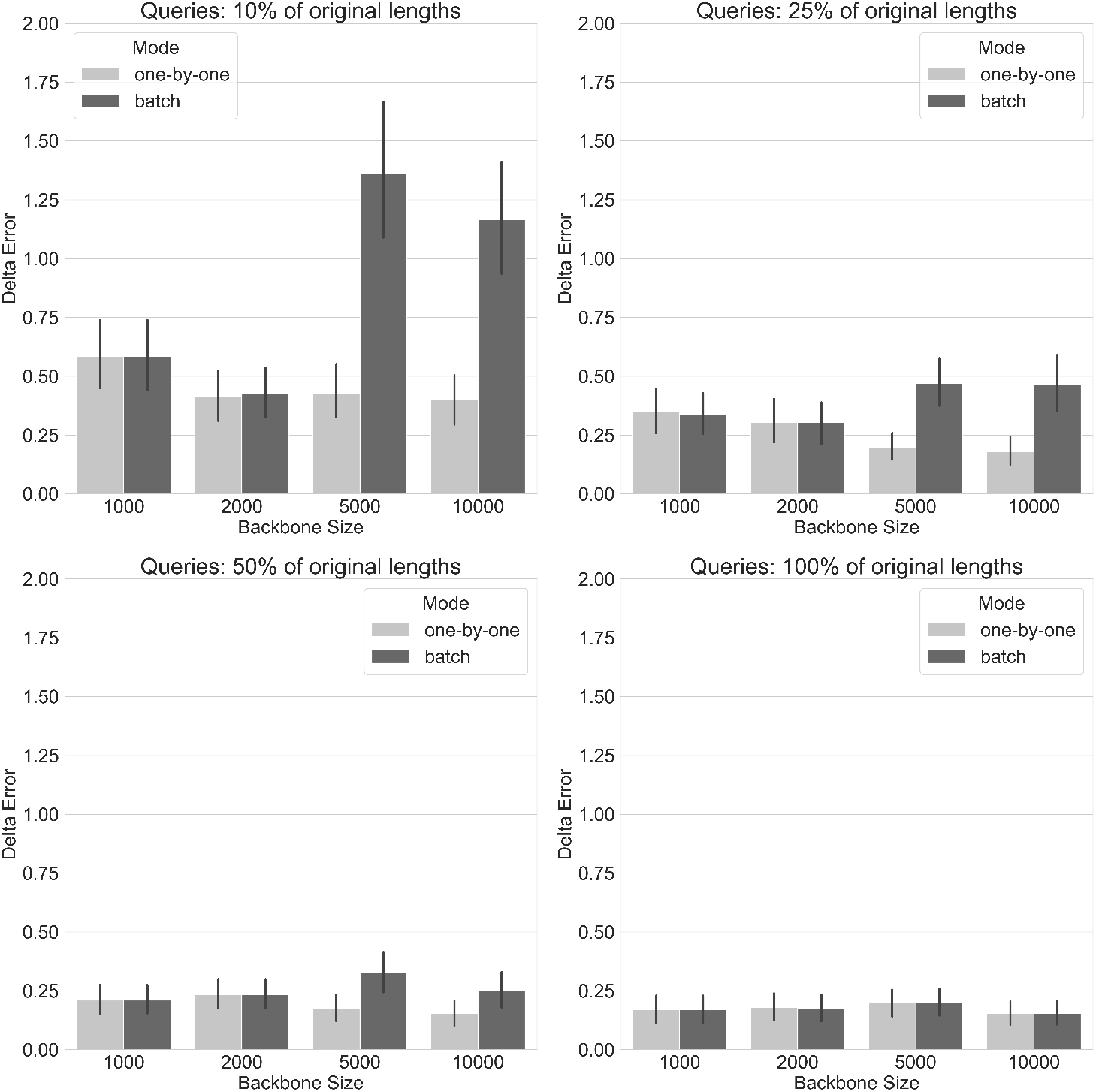
Experiment 0: Mean Delta Error of EPA-ng when placing 200 queries into a backbone tree with 1000, 2000, 5000, and 10000 leaves of the training RNASim dataset. The queries and backbone sequences are selected from the same clade. From left to right, top to bottom: query sequences (for placement) are made into fragments of 10%, 25%, 50%, and 100% of their original lengths (100% means full-length). We show two strategies for using EPA-ng for placement: “batch” means that all queries are placed at once, while “one-by-one” means that each query sequence is placed individually.

### S2 Additional Results for Experiment 1

#### S2.1 BSCAMPP Design

The SCAMPP framework scales EPA-ng to larger backbone trees, but has a linearly increasing runtime when increasing the number of query sequences. The current approach of running EPA-ng for a single query sequence on a backbone tree does not take advantage of EPA-ng’s sub-linear runtime scalability with respect to the number of query sequences.

In order to enable SCAMPP for batch query sequence processing, four new approaches are explored to enable the phylogenetic placement method to place multiple query sequences into a subtree. These four new approaches are implemented and tested specifically with EPA-ng, since this is optimized already for fast parallel placement of many query sequences.

There are four general strategies taken and tested in this study described here as EPA-ng-BSCAMPP-1, EPA-ng-BSCAMPP-2 and EPA-ng-BSCAMPP-3, and EPA-ng-BSCAMPP-4. All four approaches use a backbone tree, leaf voting strategy to inform the subtree selection process. The subtree size, *B*, is parameterized and the number of votes per query sequence, *v*, is parameterized and determined by the *v* leaves in the tree with the smallest Hamming distance to the query sequence. Once all leaves in the backbone tree have been voted on, the leaf, *l*, with the most votes is selected, and a subtree is built by greedily adding leaves with the shortest distance to *l*, until *B* leaves are in the subtree. It is at this point that the four methods differ, but in all approaches subtrees are selected, query sequences are assigned, and finally, once all queries are included in a subtree, EPA-ng is run for each subtree, query set pairing.

The four approaches are described below in more depth with their differences in bold.

##### EPA-ng-BSCAMPP-1

- The input is a set of query sequences, backbone tree for placement, and a joint backbone and query multiple sequence alignment.
- The query sequences vote on *v* leaves based on the *v* closest leaves by Hamming distance in the backbone tree.
- Then while there are query sequences not assigned to a subtree, choose the most voted leaf in the tree, build a subtree using a breadth first search collecting nearby leaves from the full tree until *B* leaves are in the subtree. Assign queries to the subtree if **they have any votes within that subtree**. And remove the votes for assigned query sequences.
- Run EPA-ng for the subtrees and their assigned query sequences.
- Determine each query sequence’s placement in the full backbone tree, and return a file containing all placements, with their requisite confidence score, distal length, placement edge number etc.

##### EPA-ng-BSCAMPP-2

- The input is a set of query sequences, backbone tree for placement, and a joint backbone and query multiple sequence alignment.
- Each query sequence votes for *v* leaves based on the *v* closest leaves by Hamming distance in the backbone tree.
- Then while there are query sequences not assigned to a subtree, choose the most voted leaf in the tree, build a subtree using a breadth first search collecting nearby leaves from the full tree until *B* leaves are in the subtree. Assign queries to the subtree if **all of their votes are contained within that subtree or if the leaf with the closest Hamming distance to the query is used to start the breadth first search**. And remove the votes for assigned query sequences.
- Run EPA-ng for the subtrees and their assigned query sequences.
- Determine each query sequence’s placement in the full backbone tree, and return a file containing all placements, with their requisite confidence score, distal length, placement edge number etc.

##### EPA-ng-BSCAMPP-3

- The input is a set of query sequences, backbone tree for placement, and a joint backbone and query multiple sequence alignment.
- Each query sequence votes for *v* leaves based on the *v* closest leaves by Hamming distance in the backbone tree.
- Let set *A* = ∅.
- Then while there are query sequences not **in set A**, choose the most voted leaf in the tree, build a subtree using a breadth first search collecting nearby leaves from the full tree until *B* leaves are in the subtree. **Add** queries to **set A** if **all of their votes are contained within that subtree or if the leaf with the closest Hamming distance to the query is used to start the breadth first search**. And remove the votes for **query sequences in set A**.
- **Assign each query sequence to the subtree with minimum total Hamming distance from the query sequence to all of the leaves in the subtree**.
- Run EPA-ng for the subtrees and their assigned query sequences.
- Determine each query sequence’s placement in the full backbone tree, and return a file containing all placements, with their requisite confidence score, distal length, placement edge number etc.

##### EPA-ng-BSCAMPP-4

- The input is a set of query sequences, backbone tree for placement, and a joint backbone and query multiple sequence alignment.
- Each query sequence votes for *v* leaves based on the *v* closest leaves by Hamming distance in the backbone tree. **Note the top vote leaf for each query as the single leaf with the smallest Hamming distance in the backbone tree**.
- Let set *A* = ∅.
- Then while there are query sequences not **in set A**, choose the most voted leaf in the tree. Let this leaf be a seed sequence, and build a subtree using a breadth first search collecting nearby leaves from the full tree until *B* leaves are in the subtree. **Add** queries to **set A** if **the top vote leaf for that query is in the subtree** And remove the votes for **query sequences in set A**.
- **Assign each query sequence to the subtree with smallest topological distance from the top voted leaf to the subtree’s seed sequence**.
- Run EPA-ng for the subtrees and their assigned query sequences.
- Determine each query sequence’s placement in the full backbone tree, and return a file containing all placements, with their requisite confidence score, distal length, placement edge number etc.

The difference between methods occurs in the third and fourth steps where the subtrees are selected and queries are assigned to subtrees. Versions 1 and 2 assign queries while building subtrees, but they do so differently. Version 1 allows a query to be assigned to the first subtree built that contains any of the *v* voted leaves with low Hamming distance. Version 2 is more selective in the query assignment, in that it will only allow a query to be used in a tree if all of its votes are contained in the tree, or if the best vote with the lowest Hamming distance is used as the leaf to start the greedy search to build the tree.

Version 3 uses the same subtrees as version 2, but it reassigns queries rather than assigning the query to the first subtree for which it may be used. The reassignment metric is as follows: each query is assigned to the subtree which minimizes the sum of the Hamming distance of the query to each leaf in the subtree.

Finally, version 4 deviates from version 3 with two adjustments. It requires the nearest leaf to be in the subtree; it does not look at other votes. It also reassigns queries to subtrees by choosing the subtree minimizing edge length distance from the nearest leaf to the subtree’s seed leaf.

**Figure S3:**
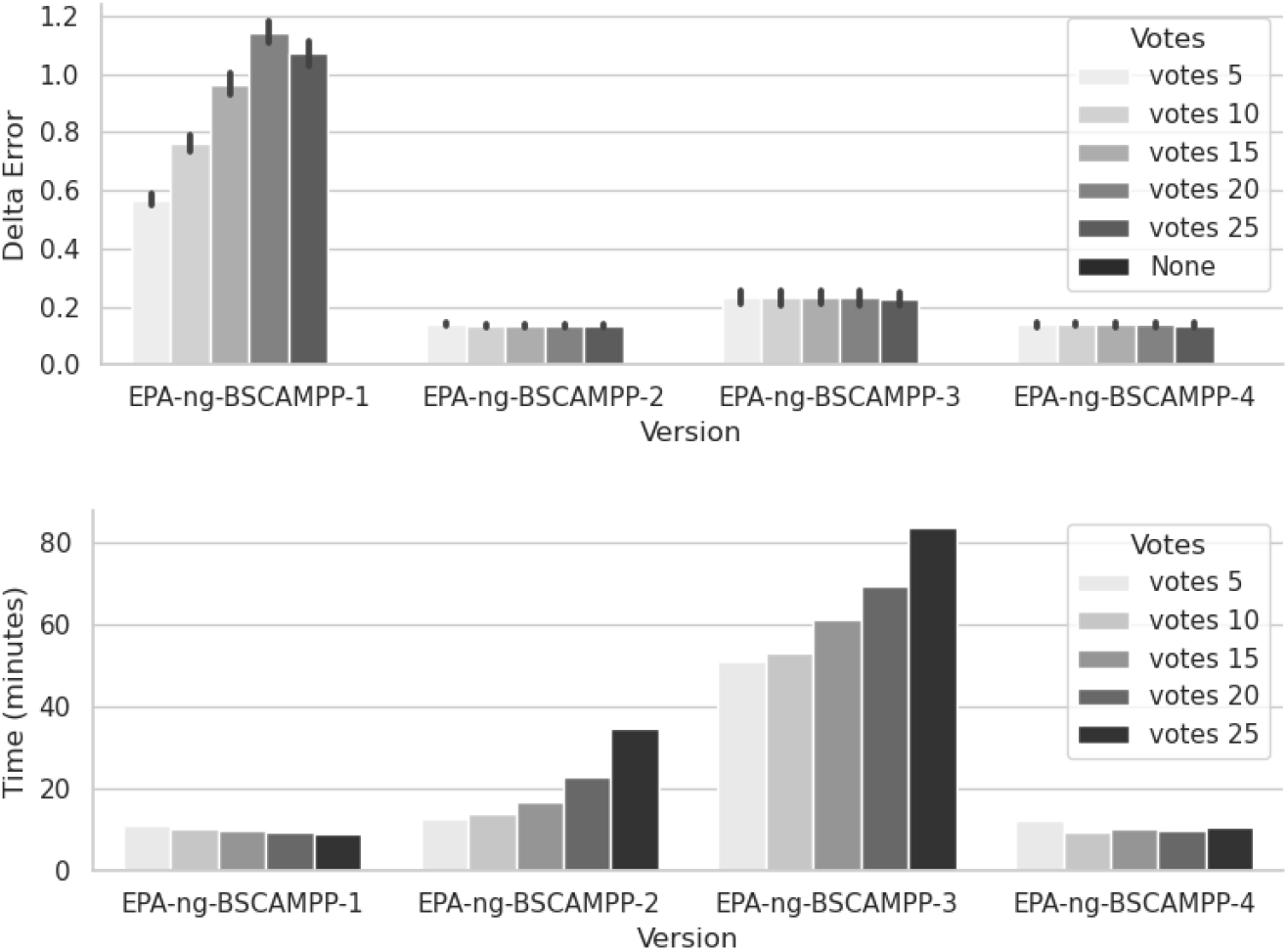
Experiment 1 (BSCAMPP Design). Top: Mean Delta Error plus standard error for placement of full length sequences on the RNASim backbone trees with 50,000 leaves and 10,000 query sequences. We show placement delta error for four versions of EPA-ng-BSCAMPP varying parameter v (the number of votes per query). Bottom: Total runtime for placement of all 10,000 full length query sequences on the RNASim backbone trees with 50,000 leaves. We show placement time for four versions of EPA-ng-BSCAMPP varying parameter v (the number of votes per query).

We test all four versions of EPA-ng-BSCAMPP on two datasets, nt78 and RNAsim. The delta error (Figure S3, top) shows that versions 2 and 4 are the most accurate. In addition, the runtime for version 4 (Figure S3, bottom) shows that EPA-ng-BSCAMPP-4 is able to place all 10,000 query sequences in just under 10 minutes when there are 10 votes, and just over 12 minutes in the worst case when there are 5 votes. EPA-ng-BSCAMPP-4 is faster than EPA-ng-BSCAMPP-2 in all cases.

EPA-ng-BSCAMPP-1 was able to place all 10,000 query sequences in around 10 minutes in all cases, but was the worst performing with respect to delta error, showing 3 to 5 times more error than EPA-ng-BSCAMPP-2 depending on the number of votes. EPA-ng-BSCAMPP-3 showed lower error than EPA-ng-BSCAMPP-1; however, it is not as accurate as EPA-ng-BSCAMPP-2 and EPA-ng-BSCAMPP-4, and takes between 50 and 80 minutes to run, giving EPA-ng-BSCAMPP-4 the advantage.

The accuracy with respect to the number of votes shows different trends based on the version. While the number of votes increases error for EPA-ng-BSCAMPP-1, it only has a small impact on the accuracy of EPA-ng-BSCAMPP-2, EPA-ng-BSCAMPP-3, and EPA-ng-BSCAMPP-4.

**Figure S4:**
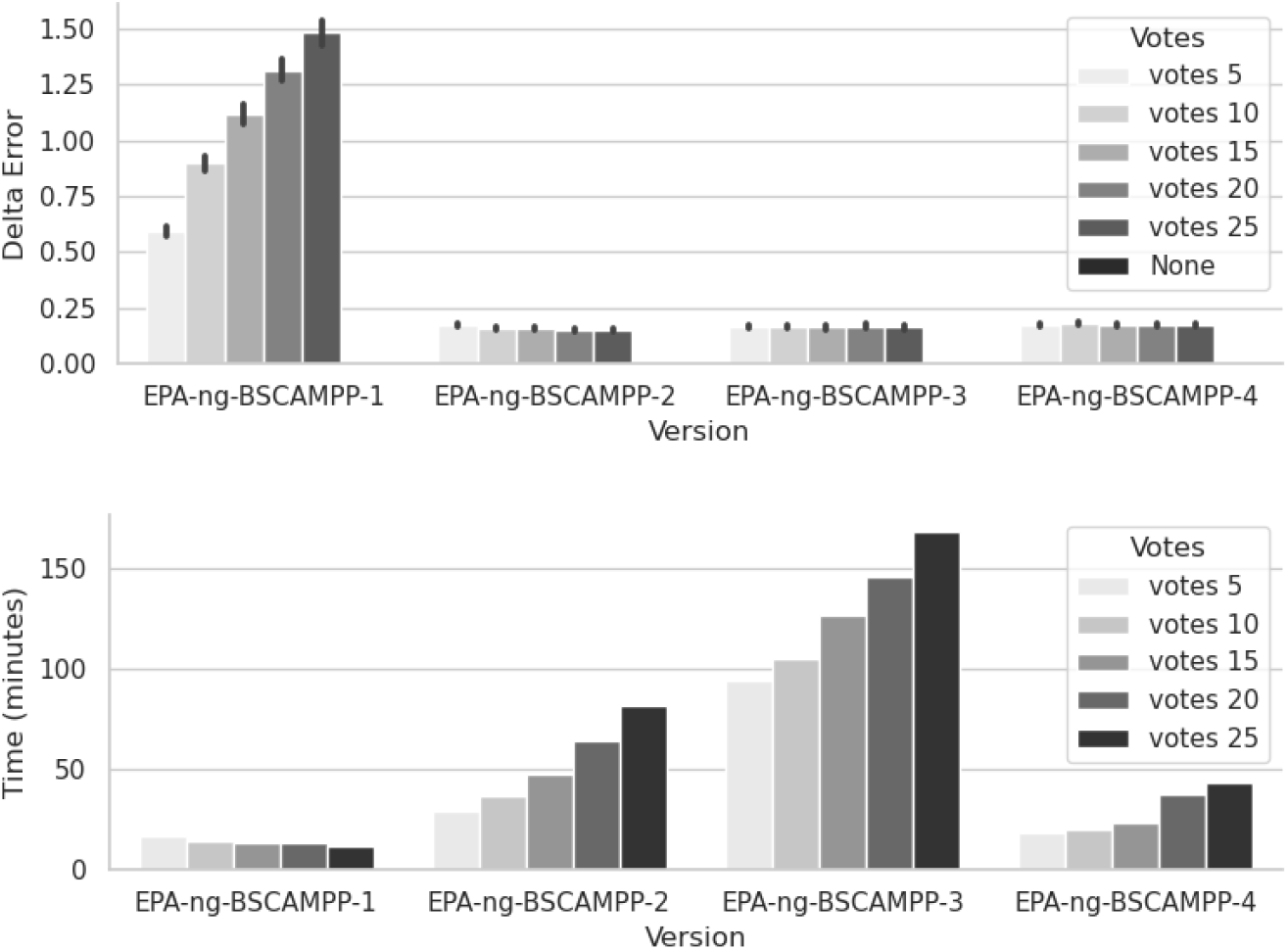
Experiment 1 (BSCAMPP Design). Top row: Mean Delta Error plus standard error for placement of full length sequences on the nt78 backbone trees with 68,132 leaves and 10,000 query sequences. We show placement delta error for four versions of EPA-ng-BSCAMPP varying parameter v (the number of votes per query). Bottom Row: Total runtime for placement of all 10,000 full length query sequences on the nt78 backbone trees with 68,132 leaves. We show placement time for four versions of EPA-ng-BSCAMPP varying parameter v (the number of votes per query).

The results from the nt78 datasets show similar trends with a few exceptions (see Figure S4). One primary difference is that on this dataset EPA-ng-BSCAMPP-3 is now showing similar accuracy to EPA-ng-BSCAMPP-2 and EPA-ng-BSCAMPP-4.

EPA-ng-BSCAMPP-1 is able to complete all placements in under 20 minutes. Of the three more accurate methods, EPA-ng-BSCAMPP-4 is the fastest, placing all 10,000 query sequences in between 18 and 45 minutes, whereas EPA-ng-BSCAMPP-2 completes all placements in between 20 and 80 minutes, with the time increasing with the number of votes. In addition, EPA-ng-BSCAMPP-3 is the slowest version tested, with runtimes between 80 and 160 minutes, again increasing with the number of votes.

EPA-ng-BSCAMPP version 4 is henceforth referred to as BSCAMPP(e), since the results indicate that this is the most efficient implementation with low delta error.

#### S2.2 Setting parameters for Version 4

**Table S1:**
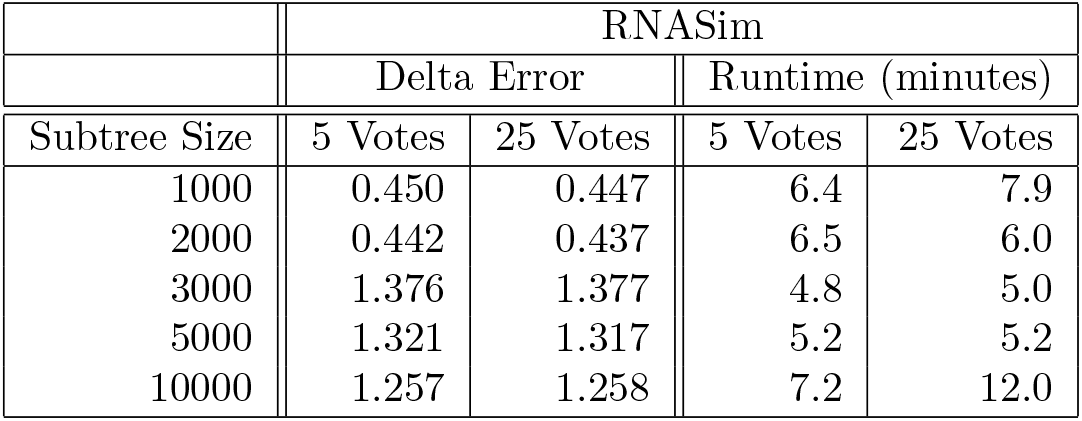
Experiment 1: Training Data Results for BSCAMPP(e) Parameter Settings on RNASim (50,000 sequences in the backbone tree with 10,000 fragmentary query sequences).

**Table S2:** Experiment 1: Testing Data Results for BSCAMPP(e) Parameter Settings on nt78 (68,132 sequences in the backbone tree with 10,000 fragmentary query sequences).

|  | nt78 |  |  |  |
| --- | --- | --- | --- | --- |
|  | Delta Error |  | Runtime (minutes) |  |
| Subtree Size | 5 Votes | 25 Votes | 5 Votes | 25 Votes |
| 1000 | 1.764 | 1.772 | 14.1 | 52.7 |
| 2000 | 1.744 | 1.734 | 14.4 | 28.4 |
| 3000 | 4.998 | 4.991 | 17.9 | 18.7 |
| 5000 | 5.028 | 5.038 | 15.5 | 18.0 |
| 10000 | 4.957 | 4.950 | 22.6 | 17.4 |

### S3 Pacbio and Illumina placed read counts for 5000M2

We report the number of reads included in the delta error computation for Experiment 4 in the main paper. APPLES-2 did not return placements for certain reads on both the Illumina and Pacbio style reads. Given this we removed them for all methods in our reporting. Figure S5 shows the placement for all methods except APPLES-2 on all 10,000 query sequences. We note that all methods delta error slightly increases when the sequences not placed by APPLES-2 are included.

**Table S3:** Experiment 4: APPLES-2 did not place certain query sequences for both Pacbio- and Illumina-style reads on the datasets with high rates of evolution. We report the number of query sequences placed out of 10,000 by APPLES-2 that are shown in the figure from the main paper for these datasets.

| Dataset | Placed Read Count |  |  |  |
| --- | --- | --- | --- | --- |
|  | Illumina |  | PacBio |  |
|  | R0 | R1 | R0 | R1 |
| 5000M2 | 9993 | 9958 | 9792 | 9909 |
| 5000M3 | 9993 | 10000 | 9998 | 9996 |
| 5000M4 | 9997 | 9998 | 10000 | 9997 |

**Figure S5:**
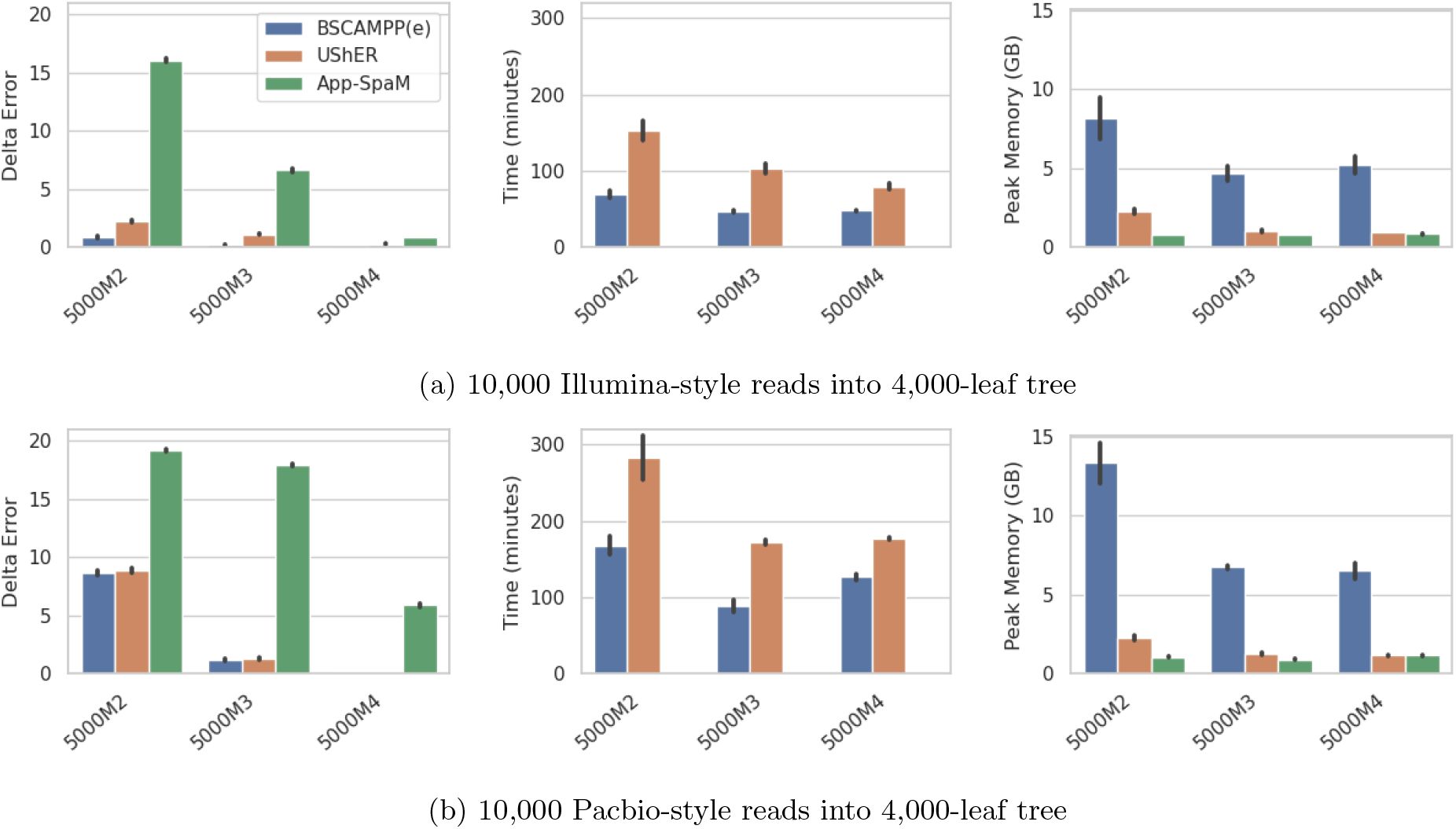
5000M2-4: Three simulated datasets with varying rates of evolution from higher (5000M2) to lower (5000M4). Shows placement of 10,000 queries into 4,000-leaf trees over two replicates for 5000M2, 5000M3, and 5000M4 (In this figure we show placement of only methods that place all 10,000 queries). From left to right: Mean Delta Error, total runtime, and peak memory usage for phylogenetic placement methods (BSCAMPP(e), UShER, and App-SpaM) on three testing datasets using **Illumina** style sequencing reads with sequencing error (above) and **Pacbio** style sequencing error (below). For BSCAMPP(e), and UShER the reads are first aligned to the reference alignment using UPP, and the aligned queries are placed into the reference alignment and tree.

### S4 Datasets and Commands

All code is available on GitHub at https://github.com/ewedell/BSCAMPP_code.

The datasets were built using the following commands, with scripts included on the GitHub site.

For the RNASim dataset, all steps are followed. For the nt78 dataset 10,000 queries were randomly selected and bullets 1 and 2 below were skipped.

In addition the 16S.B.ALL dataset contained duplicate sequences. We removed these from the reference and “true topological” tree and alignments before running any method.

1. To generate replicates, choose query sequences, and remove sites with over 95% gaps:

~~~
      python3 ./build_dataset.py -m ./true.fasta -t ./true.tt -o ./ -x a
~~~

2. To convert from RNA to DNA:

~~~
      sed ‘/^[^>]/ y/uU/tT/’ aln.fa > aln_dna.fa
~~~

3. To separate query and reference alignments:

~~~
      python3 ./build_dataset.py -m aln_dna.fa -o ./ -x b -q query.txt
~~~

4. To estimate the backbone tree before queries are pruned using FastTree v2.1.11 single precision [6]:

~~~
      FastTreeMP -nosupport -gtr -gamma -nt -log ./true.fasttree.log
      < ./aln_dna.fa > ./true.fasttree
~~~

5. For 16S.B.ALL where the published reference topology was used, we re-estimated the branch lengths for pplacer with FastTree v2.1.11 using the following command:

~~~
      FastTreeMP -nosupport -gtr -gamma -nt -mllen -nome -intree reference.tree -log true.fasttree.log
      < aln_dna.fa > true.fasttree
~~~

6. To re-estimate branch lengths for use with EPA-ng using RAxML-ng v1.0.3 [3]:

~~~
      /raxml-ng/bin/raxml-ng --evaluate --msa ref.fa --model GTR+G --tree tree
      --brlen scaled
~~~

7. To build fragmentary sequences:

~~~
      python3 get_frag_aln.py -i query.fa -p 1 -a .10 -s 10 -o a_higher_frag.fa
      -of ua_higher_frag.fa
~~~

8. To build Illumina style reads [2]:

~~~
      art_illumina -ss HS25 -sam -i [reference sequences] -l 150 -c 20
      -o [output prefix]
~~~

9. To build Pacbio style reads [5]:

~~~
      pbsim [reference sequences] --prefix [output directory] --depth 10
      --length-min 50 --length-mean 450 --accuracy-mean 0.78 --accuracy-sd 0.07
      --seed [seed number] --model_qc [CLR model path]
~~~

10. To align query sequences to reference using UPP [4] for 5000M2-M4:

~~~
      run_upp.py -x 16 -t ${dir}/backbone_epa.tree -a ${dir}/ref.fa -s ${dir}/illumina.fasta
      -m dna -A 10 -p ${dir}/ -rt -d ${dir}/ -o est.aln.illumina
~~~

11. To align query sequences to reference using UPP [4] for all other datasets with lower rates of evolution:

~~~
      run_upp.py -x 16 -t ${dir}/backbone_epa.tree -a ${dir}/ref.fa -s ${dir}/illumina.fasta
      -m dna -A 100 -p ${dir}/ -rt -d ${dir}/ -o est.aln.illumina
~~~

12. To re-estimate branch lengths under Minimum Evolution for use with APPLES-2 using FastTree v2.1.11 single precision [6]:

~~~
      FastTreeMP -nosupport -nt -nome -noml -log fasttree.log -intree reference.nwk
      < alignment.fasta > fasttree.nwk
~~~

13. To build a reference package with taxtastic taxit [1] for use with SCAMPP(p) (backbone pplacer.tree is the tree with query sequences pruned preserving branch lengths from the fasttree-2 estimated tree and true.fasttree.log is the log file created from running bullet 4 or 5 above):

~~~
      taxtastic-env/bin/taxit create -P my.refpkg -l 16S --aln-fasta ref.fa
      --tree-file backbone_pplacer.tree --tree-stats true.fasttree.log
~~~

## S5 Results Tables

We have included tables with exact results for both true and estimated alignments of query sequences on both fragmentary and full-length sequences in this section. For each dataset the mean delta error, runtime, and peak memory usage are listed.

We have not included the alignment-free methods with our results using the true query sequence alignment to the reference because the alignment-based methods would not have alignment time included, and thus a comparison would not be representative of the true time required.

### S5.1 Fragmentary Sequence Results

**Table S4:** Method comparison on all datasets using the true fragmentary query sequence alignment (backbone size is listed below each dataset). All methods are run with their default settings (BSCAMPP: *B* = 2000 and *v* = 5, SCAMPP: *B* = 2000). All query sequences are fragmentary (∼150 nt). An **X** indicates that the analysis failed due to out-of-memory issues.

| Method | RNAsim (training) - 50,000 backbone, 10,000 queries |  |  |
| --- | --- | --- | --- |
|  | Mean Delta Error | Runtime (minutes) | Peak Memory (GB) |
| BSCAMPP(e) | 0.45 | 6.1 | 3.0 |
| SCAMPP(e) | 0.43 | 457.4 | 1.2 |
| SCAMPP(p) | 0.39 | 1635.7 | 0.2 |
| EPA-ng | X | X | X |
| APPLES-2 | 1.77 | 6.2 | 0.3 |
| UShER | 0.56 | 509.9 | 1.4 |
| Method | nt78 (testing) - 68,134 backbone, 10,000 queries |  |  |
|  | Mean Delta Error | Runtime (minutes) | Peak Memory (GB) |
| BSCAMPP(e) | 1.74 | 13.7 | 2.6 |
| SCAMPP(e) | 1.72 | 429.2 | 1.2 |
| SCAMPP(p) | 1.69 | 1426.4 | 0.2 |
| EPA-ng | X | X | X |
| APPLES-2 | 3.06 | 4.1 | 0.4 |
| UShER | 1.94 | 275.5 | 0.7 |
| Method | RNAsim (testing) - 50,000 backbone, 10,000 queries |  |  |
|  | Mean Delta Error | Runtime (minutes) | Peak Memory (GB) |
| BSCAMPP(e) | 0.51 | 6.9 | 3.0 |
| SCAMPP(e) | 0.51 | 486.0 | 1.2 |
| SCAMPP(p) | 0.46 | 1421.3 | 0.2 |
| EPA-ng | X | X | X |
| APPLES-2 | 1.52 | 4.8 | 1.1 |
| UShER | 0.61 | 443.7 | 1.5 |
| Method | 16S.B.ALL (testing) - 20,000 backbone, 5,093 queries |  |  |
|  | Mean Delta Error | Runtime (minutes) | Peak Memory (GB) |
| BSCAMPP(e) | 0.58 | 3.8 | 3.0 |
| SCAMPP(e) | 0.58 | 214.6 | 1.2 |
| SCAMPP(p) | 0.57 | 952.3 | 0.2 |
| EPA-ng | 0.71 | 17.4 | 37.3 |
| APPLES-2 | 1.11 | 9.3 | 1.4 |
| UShER | 0.56 | 40.6 | 0.16 |

**Table S5:** Method comparison on two datasets using the estimated fragmentary query sequence alignment estimated using UPP (alignment time included in alignment based methods). All methods are run with their default settings (BSCAMPP: *B* = 2000 and *v* = 5, SCAMPP: *B* = 2000). All query sequences are fragmentary (∼150 nt). An **X** indicates that the analysis failed due to out-of-memory issues.

| Method | RNAsim (training) - 50,000 backbone, 10,000 queries |  |  |
| --- | --- | --- | --- |
|  | Mean Delta Error | Runtime (minutes) | Peak Memory (GB) |
| Alignment-based |  |  |  |
| BSCAMPP(e) | 0.46 | 121.0 | 3.0 |
| SCAMPP(e) | 0.45 | 546.8 | 1.6 |
| SCAMPP(p) | 0.41 | 1511.6 | 1.6 |
| EPA-ng | X | X | X |
| APPLES-2 | 1.81 | 120.3 | 1.6 |
| UShER | 0.59 | 771.8 | 1.6 |
| Alignment-free |  |  |  |
| RAPPAS | X | X | X |
| App-SpaM | 1.62 | 6.0 | 29.6 |
| Method | nt78 (testing) - 68,134 backbone, 10,000 queries |  |  |
|  | Mean Delta Error | Runtime (minutes) | Peak Memory (GB) |
| Alignment-based |  |  |  |
| BSCAMPP(e) | 1.78 | 121.0 | 2.6 |
| SCAMPP(e) | 1.75 | 546.8 | 2.2 |
| SCAMPP(p) | 1.72 | 1511.6 | 2.2 |
| EPA-ng | X | X | X |
| APPLES-2 | 3.12 | 120.3 | 2.2 |
| UShER | 1.98 | 323.8 | 2.2 |
| Alignment-free |  |  |  |
| RAPPAS | X | X | X |
| App-SpaM | 3.20 | 6.7 | 30.7 |

### S5.2 Full-Length Sequence Results

**Table S6:** Method comparison on all datasets using the true full-length query sequence alignment (backbone size is listed below each dataset). All methods are run with their default settings (BSCAMPP: *B* = 2000 and *v* = 5, SCAMPP: *B* = 2000). All query sequences are full length. An **X** indicates that the analysis failed due to out-of-memory issues.

| Method | RNAsim (training) - 50,000 backbone, 10,000 queries |  |  |
| --- | --- | --- | --- |
|  | Mean Delta Error | Runtime (minutes) | Peak Memory (GB) |
| BSCAMPP(e) | 0.14 | 11.0 | 3.0 |
| EPA-ng | X | X | X |
| APPLES-2 | 0.12 | 6.5 | 0.3 |
| UShER | 0.13 | 865.8 | 1.4 |
| Method | nt78 (testing) - 68,134 backbone, 10,000 queries |  |  |
|  | Mean Delta Error | Runtime (minutes) | Peak Memory (GB) |
| BSCAMPP(e) | 0.18 | 21.8 | 2.6 |
| SCAMPP(e) | 0.15 | 1413.2 | 2.6 |
| EPA-ng | X | X | X |
| APPLES-2 | 0.25 | 5.3 | 0.4 |
| UShER | 0.18 | 360.3 | 0.7 |
| Method | RNAsim (testing) - 50,000 backbone, 10,000 queries |  |  |
|  | Mean Delta Error | Runtime (minutes) | Peak Memory (GB) |
| BSCAMPP(e) | 0.15 | 8.4 | 3.0 |
| SCAMPP(e) | 0.15 | 1445.1 | 3.0 |
| EPA-ng | X | X | X |
| APPLES-2 | 1.76 | 4.2 | 0.3 |
| UShER | 0.14 | 730.8 | 1.5 |
| Method | 16S.B.ALL (testing) - 20,000 backbone, 5,093 queries |  |  |
|  | Mean Delta Error | Runtime (minutes) | Peak Memory (GB) |
| BSCAMPP(e) | 0.024 | 2.6 | 3.5 |
| SCAMPP(e) | 0.023 | 742.8 | 3.2 |
| EPA-ng | 0.062 | 13.0 | 64.4 |
| APPLES-2 | 0.257 | 4.5 | 1.4 |
| UShER | 0.063 | 32.8 | 0.1 |

**Table S7:** Method comparison on two datasets using the estimated full-length query sequence alignment, estimated using UPP (alignment time included in alignment based methods). All methods are run with their default settings (BSCAMPP: *B* = 2000 and *v* = 5, SCAMPP: *B* = 2000). An **X** indicates that the analysis failed due to out-of-memory issues.

| Method | RNAsim (training) - 50,000 backbone, 10,000 queries |  |  |
| --- | --- | --- | --- |
|  | Mean Delta Error | Runtime (minutes) | Peak Memory (GB) |
| Alignment-based |  |  |  |
| BSCAMPP(e) | 0.14 | 125.9 | 3.0 |
| EPA-ng | X | X | X |
| APPLES-2 | 0.13 | 121.6 | 1.6 |
| UShER | 0.13 | 916.7 | 1.6 |
| Alignment-free |  |  |  |
| RAPPAS | X | X | X |
| App-SpaM | 0.60 | 88.1 | 60.5 |
| Method | nt78 (testing) - 68,134 backbone, 10,000 queries |  |  |
|  | Mean Delta Error | Runtime (minutes) | Peak Memory (GB) |
| Alignment-based |  |  |  |
| BSCAMPP(e) | 0.18 | 76.2 | 2.6 |
| EPA-ng | X | X | X |
| APPLES-2 | 0.28 | 59.8 | 2.2 |
| UShER | 0.18 | 391.0 | 2.2 |
| Alignment-free |  |  |  |
| RAPPAS | X | X | X |
| App-SpaM | X | X | X |

#### S5.3 Tables on ultra-large references and datasets with high rates of evolution

**Table S8:**
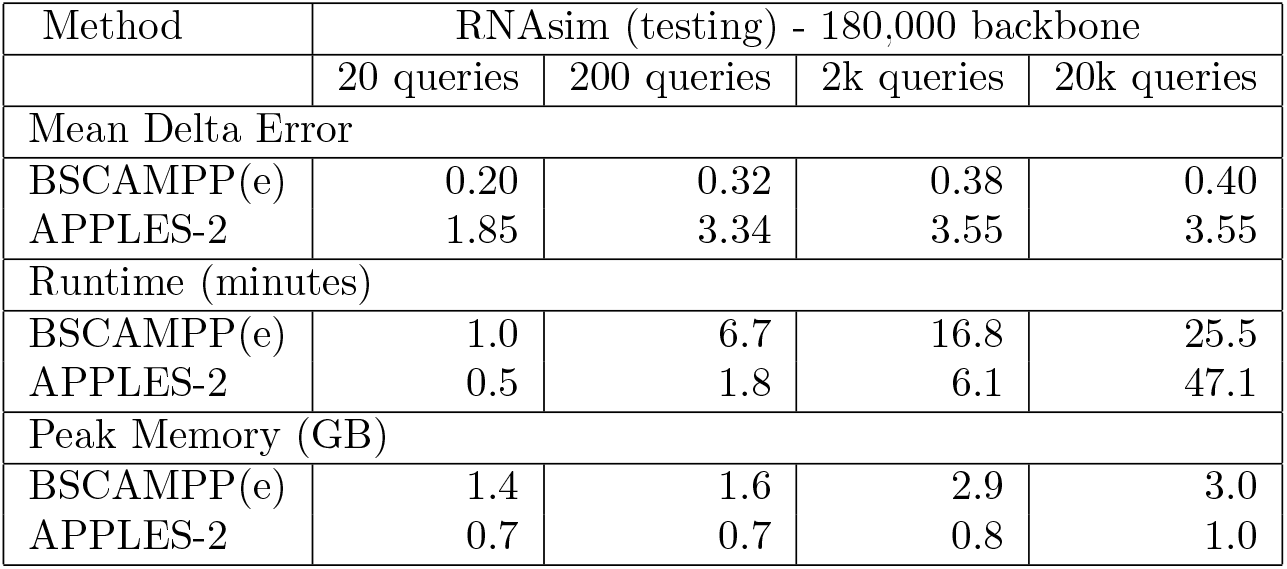
Method comparison on 180K leaf tree varying query sequence set size, using the true fragmentary query sequence alignment to the reference. Fragmentary sequences are generated without sequencing error. Both methods are run with their default settings (BSCAMPP: *B* = 2000 and *v* = 5). All query sequences are fragmentary (∼150 nt).

**Table S9:** Method comparison on testing datasets with varying rate of evolution using the true fragmentary query sequence alignment to the reference. Fragmentary sequences are generated without sequencing error. Both methods are run with their default settings (BSCAMPP: *B* = 2000 and *v* = 5). Fragmentary sequences are 10% their original length.

| Method | 5000M2-4 (testing) - 4,000 backbone, 10,000 fragmentary queries |  |  |
| --- | --- | --- | --- |
|  | Mean Delta Error | Runtime (minutes) | Peak Memory (GB) |
| 5000M2 |  |  |  |
| BSCAMPP(e) | 1.37 | 1.4 | 8.0 |
| UShER | 2.83 | 80.8 | 0.3 |
| 5000M3 |  |  |  |
| BSCAMPP(e) | 0.47 | 0.8 | 4.4 |
| UShER | 0.77 | 55.2 | 0.3 |
| 5000M4 |  |  |  |
| BSCAMPP(e) | 0.36 | 0.7 | 5.4 |
| UShER | 0.73 | 32.8 | 0.2 |

